# Microbiota-specific serum IgG links gut and joints through immune–endothelial crosstalk in arthritis

**DOI:** 10.64898/2026.01.29.702529

**Authors:** Eva Schmid, Nadine Otterbein, Stephen Ariyeloye, Heike Danzer, Michael Frech, Elisabeth Naschberger, Marco Munoz Becerra, Kerstin Sarter, Georg Schett, Alf Kastbom, Anna Svärd, Ben Wielockx, Mario M. Zaiss

**Affiliations:** Department of Internal Medicine 3, Rheumatology and Immunology, Friedrich-Alexander-Universität Erlangen-Nürnberg (FAU) and Universitätsklinikum Erlangen, Erlangen, Germany; Deutsches Zentrum Immuntherapie (DZI), Friedrich-Alexander-Universität Erlangen-Nürnberg (FAU) and Universitätsklinikum Erlangen, Erlangen, Germany; Institute of Clinical Chemistry and Laboratory Medicine, Technische Universität Dresden, 01307, Dresden, Germany; Experimental Centre, Faculty of Medicine, Technische Universität Dresden, 01307, Dresden, Germany; Division of Molecular and Experimental Surgery, Uniklinikum Erlangen, Friedrich-Alexander-Universität (FAU) Erlangen-Nürnberg, Erlangen, Germany; Linköping University, 58183 Linköping, Sweden; Falun Hospital, 79182 Falun, Sweden

**Keywords:** rheumatoid arthritis, microbiota specific IgG, endothelial cells, gut, bone

## Abstract

Rheumatoid arthritis (RA) pathogenesis involves early gut immune alterations that precede clinical onset and systemic bone involvement. Using mouse and human imaging mass cytometry (IMC) and tissue sequencing, this study shows that intestinal endothelial and immune changes emerge before or coincide with arthritis symptom development. In the collagen-induced arthritis (CIA) model, intestinal vascular permeability and endothelial gene activation promoting leukocyte trafficking appeared prior to synovial inflammation. Spatial mapping of murine and human ileal tissues predicted enhanced epithelial–immune interactions and lymphoid activation, suggesting mucosal immune priming before joint pathology. Both gut-selective α4β7 integrin blockade with vedolizumab and endothelial barrier enhancement by imatinib significantly reduced arthritis severity in CIA mice. After clinical onset, microbiota-specific IgG responses expanded to recognize rare gut bacteria, reflecting increased microbial exposure. Bone marrow endothelium exhibited interferon-I–driven inflammation and vascular activation, indicating tissue-specific endothelial dysfunction. Microbiota-reactive IgG increased during CIA – likely a response to translocating gut bacteria and immune cell activation. Integrating mouse and human data, these findings define a mechanistic framework where endothelial barrier impairment, microbial translocation, and systemic endothelial activation initiate RA autoimmunity, revealing endothelial and mucosal pathways as targets for early intervention.

**Graphical Abstract:** 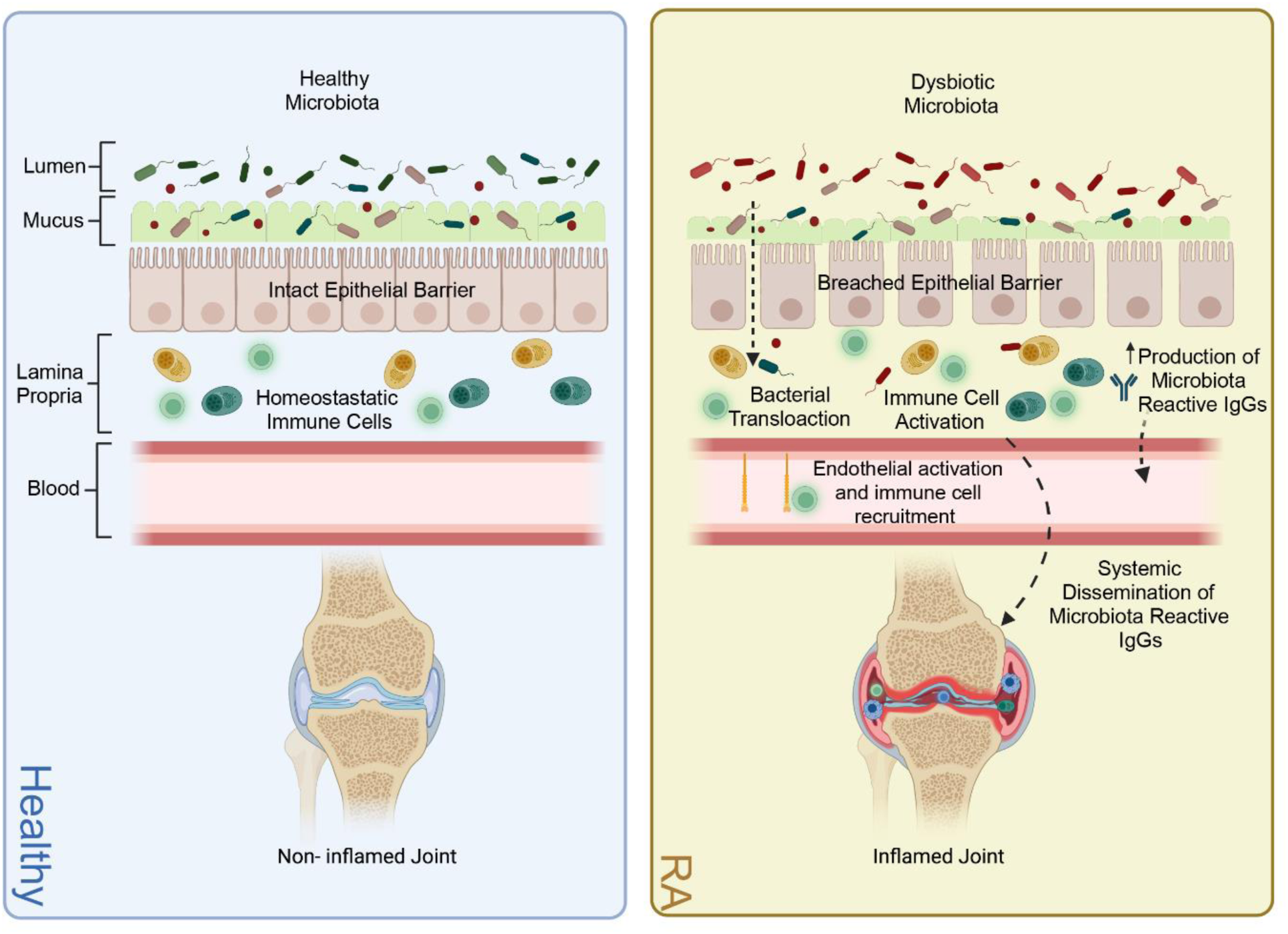

## Introduction

Rheumatoid arthritis (RA) is an autoimmune disease characterized by chronic synovial inflammation, progressive joint destruction, and systemic complications that lead to significant morbidity and reduced quality of life (1). Although genetic predisposition and environmental factors - such as smoking and microbial exposures - modulate disease susceptibility, the precise triggers initiating RA remain incompletely understood (2). Increasing evidence suggests that the intestinal tract may play a pivotal role in the early stages of RA development (3–7).

The emerging concept of inter-organ communication in RA (8) may extend beyond inflammation-mediated pathways to include direct early crosstalk between the gut and bone (9) before synovial inflammation occurs. Notably, microbiota-derived metabolites such as short-chain fatty acids (SCFAs), as well as intestinal colonization by *Prevotella* species, have been implicated in modulating bone density independent of inflammation (10, 11). Intestinal barrier leakage was shown in animal models of arthritis and RA patients (5, 7), even prior to arthritis onset (12). However, while direct bacterial translocation has not been conclusively demonstrated in RA, biomarkers indicative of microbial passage into the circulation - such as circulating bacterial DNA (16S rRNA gene copies) (13), lipopolysaccharide-binding protein (LBP), and soluble CD14 (sCD14) (14) - are elevated in patients’ blood. In autoimmune diseases such as systemic lupus erythematosus (SLE), microbial translocation is known to promote chronic inflammation and autoantibody production by activating innate immune pathways and proinflammatory cytokine release (15, 16). Persistent bacterial translocation together with local mucosal immune activation in the intestine can induce systemic microbiota-specific immunoglobulin G (IgG) (17, 18) similar to what was described following the translocation of gastrointestinal pathogens (19, 20). Although commensal gut bacteria can elicit systemic IgG under steady-state conditions (21, 22), we hypothesised that heightened systemic cellular (23) or IgG responses against the gut microbiota may act as an additional trigger for arthritis onset and directly impact systemic bone density. Systemic IgG antibodies enter the bone marrow via its highly vascularized network of endothelial cells, mainly in the sinusoids. Anti-citrullinated protein antibodies (ACPAs) are present in the bone marrow of patients with RA and have a significant role in promoting osteoclast activation and bone resorption locally within this tissue (24). Recent findings further confirmed that serum anti-modified protein antibodies (AMPAs) in RA patients bind gut microbes (25). Together, these findings highlight how local immune activation in endothelial cells in the gut and the bones and microbiota-specific IgG may jointly impact bone erosion and disease onset in RA.

We used the collagen-induced arthritis (CIA) mouse model, which recapitulates key features of preclinical RA (26). Time-course analyses showed increased intestinal endothelial permeability in the pre-disease phase, indicating early vascular dysfunction. Intestinal endothelial transcriptomes revealed heightened activation and immune-cell recruitment before clinical onset, whereas bone endothelium displayed strongest transcriptional changes during active arthritis, demonstrating tissue-specific endothelial regulation. Longitudinal microbiota profiling revealed dynamic bacterial shifts accompanied by rising microbe-specific serum IgG levels, with expanding recognition of rare taxa, indicating progressively broadened systemic immune activation similar to inflammatory bowel disease (IBD) (27). Spatial immunophenotyping of mouse and human ileal biopsies by imaging mass cytometry (IMC) together with neighbourhood analysis identified enhanced immune activation and localized cellular clusters in early RA and IBD, in contrast to the healthy-like profile of established RA, emphasizing that intestinal immune disturbances arise predominantly during early disease.

Collectively, by linking gut-derived immune responses, vascular activation, and microbiota-specific serum IgG with potential effects on bone homeostasis, this study offers mechanistic insight into early RA pathogenesis and highlights the gut as a promising target for early intervention.

## Results

### Inflammatory arthritis alters cellular interactions in the ileum

To investigate intestinal changes in arthritis, we applied an IMC panel targeting endothelial, immune, and stromal cells (Supplementary Fig. 1, Supplementary Table 1) to ileal sections from naïve mice and CIA mice in order to define the changes in the cellular landscape and the cell interactions in the gut during the onset of arthritis. Mice were sacrificed every five days post-immunization (dpi) until 40 dpi. Three mice per timepoint and three region of interests (ROIs) per mouse yielded 847,433 single cells (Supplementary Table 2). Rphenograph (k=50) identified 21 clusters, which were assigned to cell types based on marker expression and spatial location (**Figs. 1A and B**, Supplementary Fig. 2, Supplementary Table 3). Paw swelling, and synovial inflammation in CIA mice began at 25 dpi (**Fig. 1C**), but compared to naïve ileal histology, overall cell frequencies remained largely unchanged in IMC analysis in CIA mice (**Fig. 1D**). However, spatial single-cell analysis allowed estimation and prediction of cell–cell interactions between cell types using Delaunay triangulation (max dist.=20, **Fig. 2A**). Here, cellular neighbourhoods (CN) were defined based on these interaction predictions and remained stable across inflammatory arthritis in CIA mice with CN1 (rich in CD8 T cells), CN2 (rich in epithelial/shedded cells), CN3 (rich in vascular endothelial cells (VECs) /macrophages), CN4 (rich in epithelial), CN5 (rich in proliferating crypts/macrophage subset 1), and CN6 (rich in smooth muscle cells (SMC)s/nerve fibres) (**Figs. 2B and C**). Although the overall composition of CNs remained largely stable over time, the predicted interactions between specific immune cell types and other cell clusters shifted during disease progression. In the pre-disease phase, interactions between macrophage subset 2 and epithelial cells were predicted to be elevated within CN3 at 5 dpi. By 10 dpi, increased predicted interactions between macrophage subset 2 and intraepithelial lymphocytes (IELs) with VECs were observed. Following the onset of synovial inflammation, enhanced predicted interactions between CD8 T cells and epithelial cells appeared in CN3 at 35 dpi, while an increase in self-interactions among macrophage subset 1 occurred in CN5 at 40 dpi. No significant changes in immune cell–cell interactions were predicted in CN2, CN4, or CN6 (see **Fig. 2D**). Taken together, intestinal cell-cell interactions in the pre-disease phase are predicted to be markedly changed during the development of arthritis.

**Fig. 1.**
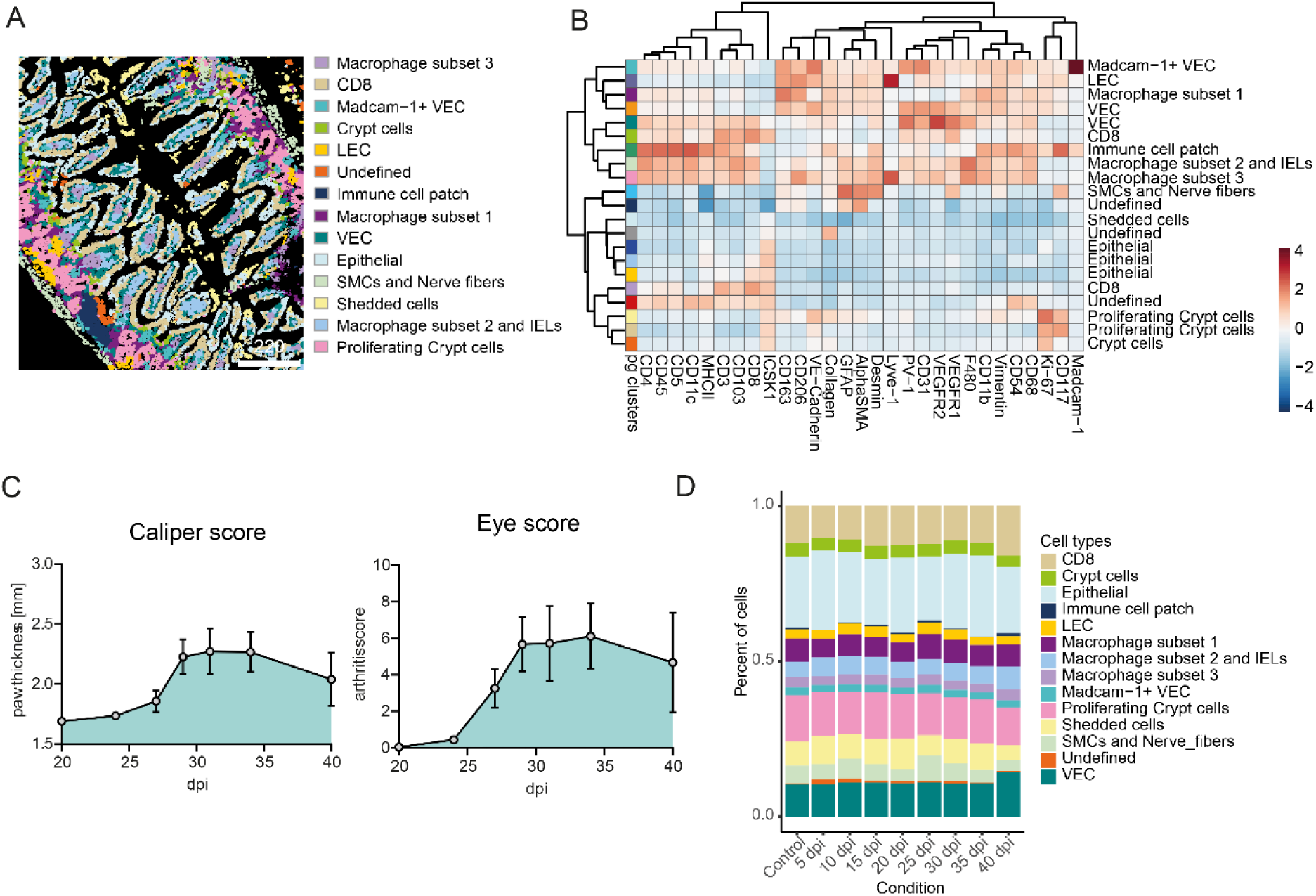
IMC analysis of ileal tissue does not reveal cellular changes during CIA. (**A**) Cell plots from one example ROI of the based-on marker expression and localization defined cell types. (**B**) Heatmap of normalized marker expression with color codes for 21 phenograph clusters and the respective defined cell types. The heatmap colors represent the z-score of the average expression of a given marker for each identified cluster. (**C**) Arthritis scores of mice after the second immunization 21 days post-immunization (dpi). Left graph shows the caliper score (mean of paw thickness +- SEM). Right graph shows the total eye score (mean +- SEM). (**D**) Bar graph showing the mean percentage of cell types across mouse samples after the different timepoints after immunization in ileal sections (n ≥ 3). Control mouse samples derived from untreated control mice, day 0 or day 40.

**Fig. 2.**
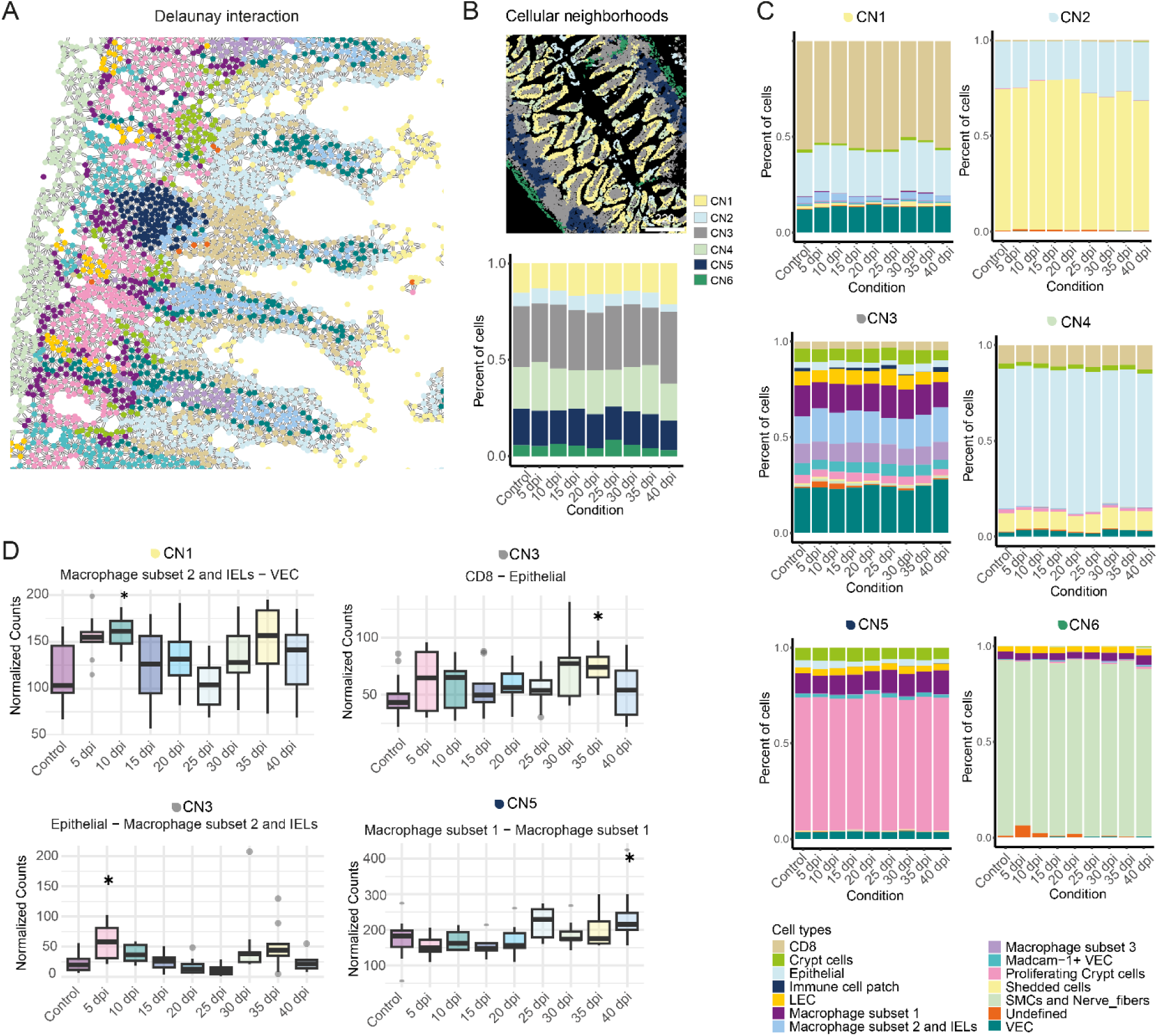
Predicted cellular interactions with immune cells are changed in the disease and pre-disease state. (**A**) Delaunay triangulation-based interaction prediction of the different cell types in one example ROI. Figure legend see lower right bottom. (**B**) Example ROI of cellular neighborhoods (CN)1 to CN6 detected in our data-set (upper part). Bar graph showing the mean percentage of cells of the different CNs at the different the dpi (lower part). (**C**) Bar graphs showing the mean percentage of cells of the different cell types in CN1 to CN6 (upper left to lower right) at the different days post immunization (dpi). (**D**) Box plots showing the normalized predicted interactions (based on Delaunay triangulation) over time of the different cell types in one CN per ROI grouped by condition. CNs are indicated above the graphs. Number of predicted interactions between cell types in one CN per ROI were normalized on the cell number in the CN per ROI (nCNinteractions/nCNcellcount *100). Only significantly different cell type immune – cell type predicted interactions to the control are shown (one-way ANOVA p<0.05 & Tukey HSD p<0.05). Data are presented as boxplots showing the median (line), interquartile range (IQR, box), and whiskers extending to 1.5× IQR. Outliers are shown as individual points. Statistical analysis was performed using one-way ANOVA. Multiple comparisons between the groups were performed with Tukey HSD. * p-value < 0.05 (n ≥ 3). Control mouse samples derived from untreated control mice, day 0 or day 40.

### Early endothelial activation in small intestine during development of arthritis

It was shown that a status of increased gut leakiness occurs in many inflammatory conditions, including RA, even before disease onset (5, 7, 12, 28). While previous studies focused on the epithelial barrier (5, 7, 29), endothelial integrity can also be compromised, as shown in celiac disease (30). To test this, we performed in vivo imaging in CIA mice at 15 dpi (pre-disease) and co-housed naïve control mice. Mice received intravenous injections of fluorescently labelled lectin to mark vessels and 70kDa dextran, which remains in intact vasculature but accumulates in crypts if the endothelial barrier is leaky. Representative images suggested an increase in FITC-Dextran-positive crypts in pre-diseased CIA mice (**Fig. 3A**). We next investigated transcriptional changes in endothelial cells, by sorting viable CD45^−^ CD31^+^ cells from the small intestine followed by bulk RNA-seq. in unimmunized mice at 0 and 50 dpi (naïve controls) and in CIA mice at 15 (pre-disease), 25 (early disease), 35 (disease), and 50 (remission) dpi (**Fig. 3B**, disease scores Supplementary Figure 9). PCA revealed a clear and progressive separation of pre-disease and early disease from healthy controls, with PC1 explaining 51% of the overall variance and therefore representing the dominant axis of transcriptional change across arthritis development. Notably, the genes contributing most strongly to PC1 were enriched for pathways involved in endothelial development and actin-filament assembly (**Figs. 3C and D**) - core biological processes influencing endothelial barrier integrity (31, 32) - highlighting their central role in distinguishing diseased from non-diseased states. Differential expression analysis revealed the strongest changes again in the pre-disease phase (n=148 genes), followed by the early disease (n=83) and disease phase (n=7); no differentially expressed genes (DEGs) were detected in the remission phase (**Fig. 3E**). Of note, the pre-disease phase was also the time when most significant cell-cell interactions were predicted by IMC as shown in Figure 1. At the pre-disease state, upregulated genes included *Sele*, *Madcam-1*, *Glycam-1* (endothelial activation and immune recruitment) and *Ackr1* (angiogenesis, inflammation), associated with leukocyte migration and adhesion (**Figs 3F and G**). During the early disease state, genes such as *Cdhr2*, *Cdhr5*, *Fabp2* were upregulated, with gene ontology (GO) terms highlighting leukocyte adhesion and receptor-mediated endocytosis (**Fig. 3H and I**). By the disease phase, *Irf7*, *Isg15*, and *Sele* were upregulated, reflecting activation of type 1 interferon (IFN) signalling (**Fig. 3J**). Overall, the intestinal endothelial RNA expression profile reveals most changes during the pre-clinical disease phase in CIA mice. In line with this observation, vedolizumab, an antibody blocking integrin-mediated intestinal homing of immune cells, as well as targeting the endothelial barrier function using Imatinib-mesylate (33) showed a tendency to ameliorate CIA (Supplementary Fig. 3).

**Fig. 3.**
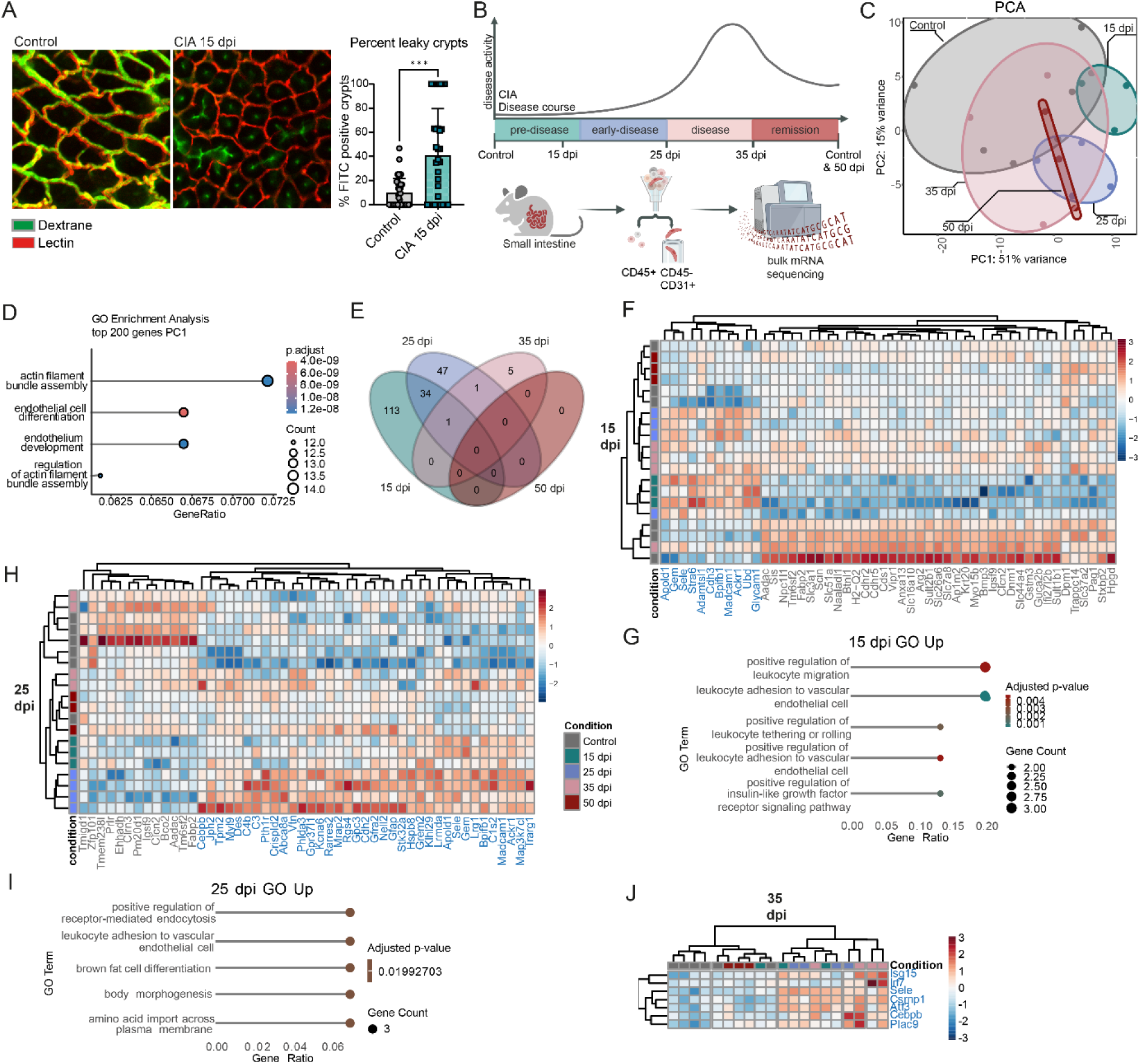
CIA changes transcriptional profile of small intestinal endothelial cells already in the preclinical phase. **(A**) Representative immunofluorescent images of colon vasculature of control mice and CIA mice 15 dpi (days post immunization) injected intravenously with FITC labelled dextran (70 kDa, measure of leakiness) and Cy5 labeled Lectin (labels blood vessels) (left side). Percentage of FITC positive crypts of total crypts (right side) per region of interest (ROI). Data are presented as mean ± SD. Statistical significance was assessed using Mann-Whitney U test. *** p-value < 0.01 (n ≥ 4, with 6 analyzed ROIs per mouse). (**B**) Graphical illustration of experimental setup. Small intestinal endothelial cells (CD45- CD31+) from control (untreated, day 0 and day 50) and CIA mice (15 dpi, 25 dpi, 35 dpi and 50 dpi) were FACS sorted and analyzed by bulk RNA sequencing. (**C**) Principal component analysis (PCA) plot of bulk RNA sequencing data from sorted endothelial cells. (**D**) GO analysis of top 200 most variable genes of PC1. (**E**) Venn diagram of differentially expressed genes (DEGs: padj < 0.05, log2FC >1.5 OR < -1.5). Differential expression analysis was performed using the DESeq2 package, with significance assessed based on p-values adjusted for multiple testing using the Benjamini-Hochberg FDR control method. (**F**) Heatmap of DESeq2 normalized gene-expression of top 50 (padj) 15 dpi DEGs. (**G**) GO analysis of 15 dpi upregulated DEGs (padj < 0.05, log2FC >1.5). Only selected terms are shown. (**H**) Heatmap of DESeq2 normalized gene-expression of top 50 (padj) 25 dpi DEGs. (**I**) GO analysis of 25 dpi upregulated DEGs (padj < 0.05, log2FC >1.5). Only selected terms are shown. (**J**) Heatmap of DESeq2 normalized gene-expression 35 dpi DEGs. Differential expression analysis was performed using the DESeq2 package, with significance assessed based on p-values adjusted for multiple testing using the Benjamini-Hochberg FDR control method. (n ≥ 3)

### Type I Interferon response in bone marrow endothelial cells of pre- arthritic mice

To assess if the transcriptional changes in endothelial cells are restricted to the intestine or if it connects the gut to the bone marrow in the pre-disease phase, we FACS-sorted viable Lineage^−^Sca1^−^ CD31^+^ cells from bone and bone marrow in naïve controls, at the pre-disease, early disease, active disease and remission state (**Fig. 4A**, disease scores Supplementary Figure 8). As an additional marker we included endomucin which is only expressed on venous and capillary endothelial cells, and a higher expression of which is linked to the inhibition of immune cell infiltration into tissue (34). At pre-disease and early disease endomucin^+^ cells were reduced compared to naïve mice, whereas no changes were observed during disease or remission phase (**Fig. 4B**). PCA plots revealed clear separation of pre-disease, early and disease phase samples from controls, with PC1 explaining 46% of the variance (**Fig. 4C**). Bulk RNA-seq. deconvolution using mMCP estimated endothelial cell frequencies to be stable across all timepoints (**Fig. 4D**), but indicated B cell contamination in sorted cells. This contamination was especially apparent in control mice and lowest in the early disease and active disease state. DEGs shared between the early disease and active disease phase, or across all groups, were enriched for B cell–related GO terms. To reduce this bias, all B cell–associated DEGs were removed from further analyses (Supplementary Table 4, Supplementary Fig. 4). Differential gene expression analysis revealed the most changes at active disease state (998 DEGs), followed by the early disease state (199 DEGs) compared to controls (**Fig. 4E**). At the pre-disease state, upregulated genes included *Oas2*, *Oas1a*, *Oasl1*, *Ifi44* and *Ifi44I* (**Fig. 4F**), type I IFN–related genes involved in positive regulation of interferon production (**Fig. 4G**). Similar GO terms were upregulated during the early disease phase (**Figs. 4H and I**). At the disease state, during active joint and bone inflammation, upregulated GO terms included positive regulation of leukocyte activation and inflammatory response (**Figs. 5A and B**). In the resolution phase, DEGs were enriched for negative regulation of inflammatory pathways and immune responses (**Figs. 5C and D**). Type I interferons can enhance endothelial activation and vascular leakage (35, 36). Consistently, *Angpt1* steadily decreased, reaching its lowest expression during the early disease phase (**Fig. 5E**), whereas *Tie1* increased during the early disease phase, promoting a pro-angiogenic phenotype via *Tie2* downregulation (**Fig. 5F**) (37). The adhesion protein *Vcam1* increased throughout disease, peaking at early disease (**Fig. 5G**), and *Selp*, expressed by endothelial cells, was elevated at early disease (**Fig. 5H**) (38).

**Fig. 4.**
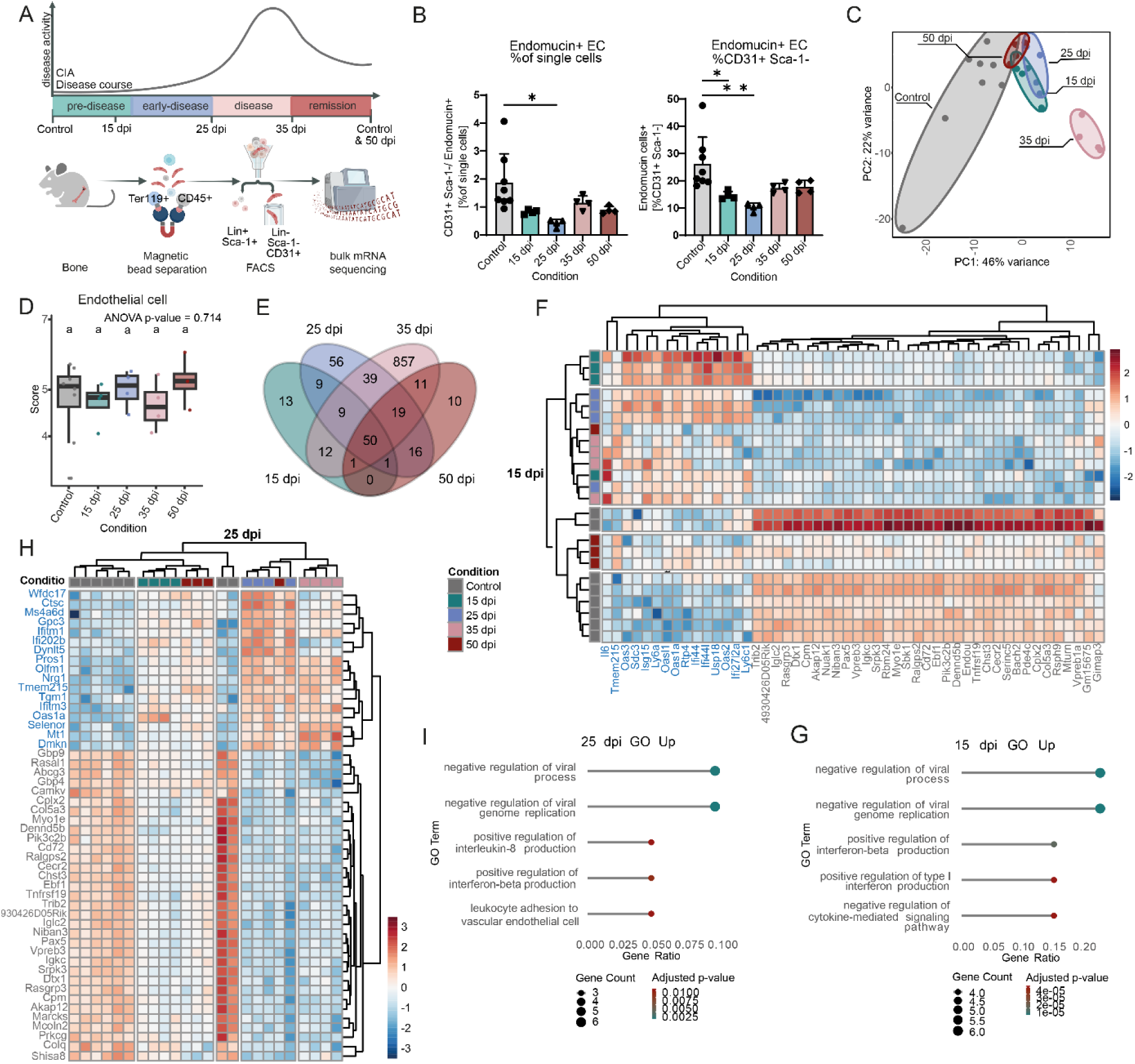
Interferon I signaling pathway in bone endothelial cells is transcriptionally induced by CIA before onset of clinical symptoms. (**A**) Graphical illustration of experimental setup. Bone and bone marrow endothelial cells (Lin- Sca1-CD31+) from Control (untreated, day 0 and day 50) and CIA mice (15, 25, 35 and 50 days post immunization (dpi)) were sorted by magnetic separation and FACS and analyzed by bulk RNA sequencing. (**B**) Percentage of endomucin^+^ endothelial cells from single cells (left side) and CD31^+^ Sca1^+^ (right side). Data are presented as mean ± SD. Statistical significance was assessed using one-way ANOVA. Multiple comparisons were performed using Tukey HSD. (n ≥ 3) * p-value < 0.05 ** p-value < 0.01. (**C**) Principal component analysis (PCA) of bulk RNA sequencing data from sorted endothelial cells. (**D**) Boxplot of estimated cell type scores (Endothelial cells) from deconvolution of bulk sequencing data using mMCP-Counter. Data are presented as boxplots showing the median (line), IGR (box), and whiskers extending to 1.5× IQR. Outliers are shown as individual points. Statistical analysis was performed using one-way ANOVA. For multiple comparisons between the groups Tukey-test was performed bp-value < 0.05.(**E**) Venn diagram of differentially expressed genes (DEGs: padj < 0.05, log2FC >1.5 OR < -1.5) using prefiltered data. B cell related genes were filtered before analysis. (**F**) Heatmap of DESeq2 normalized gene-expression of top 50 (padj) of 15 dpi DEGs. (**G**) GO analysis of 15 dpi upregulated DEGs (padj < 0.05, log2FC >1.5). (**H**) Heatmap of DESeq2 normalized gene-expression of top 50 (padj) of 25 dpi DEGs. (**I**) GO analysis of 25 dpi upregulated DEGs (padj < 0.05, log2FC >1.5). Differential expression analysis was performed using the DESeq2 package, with significance assessed based on p-values adjusted for multiple testing using the Benjamini-Hochberg FDR control method (n ≥ 3).

**Fig. 5.**
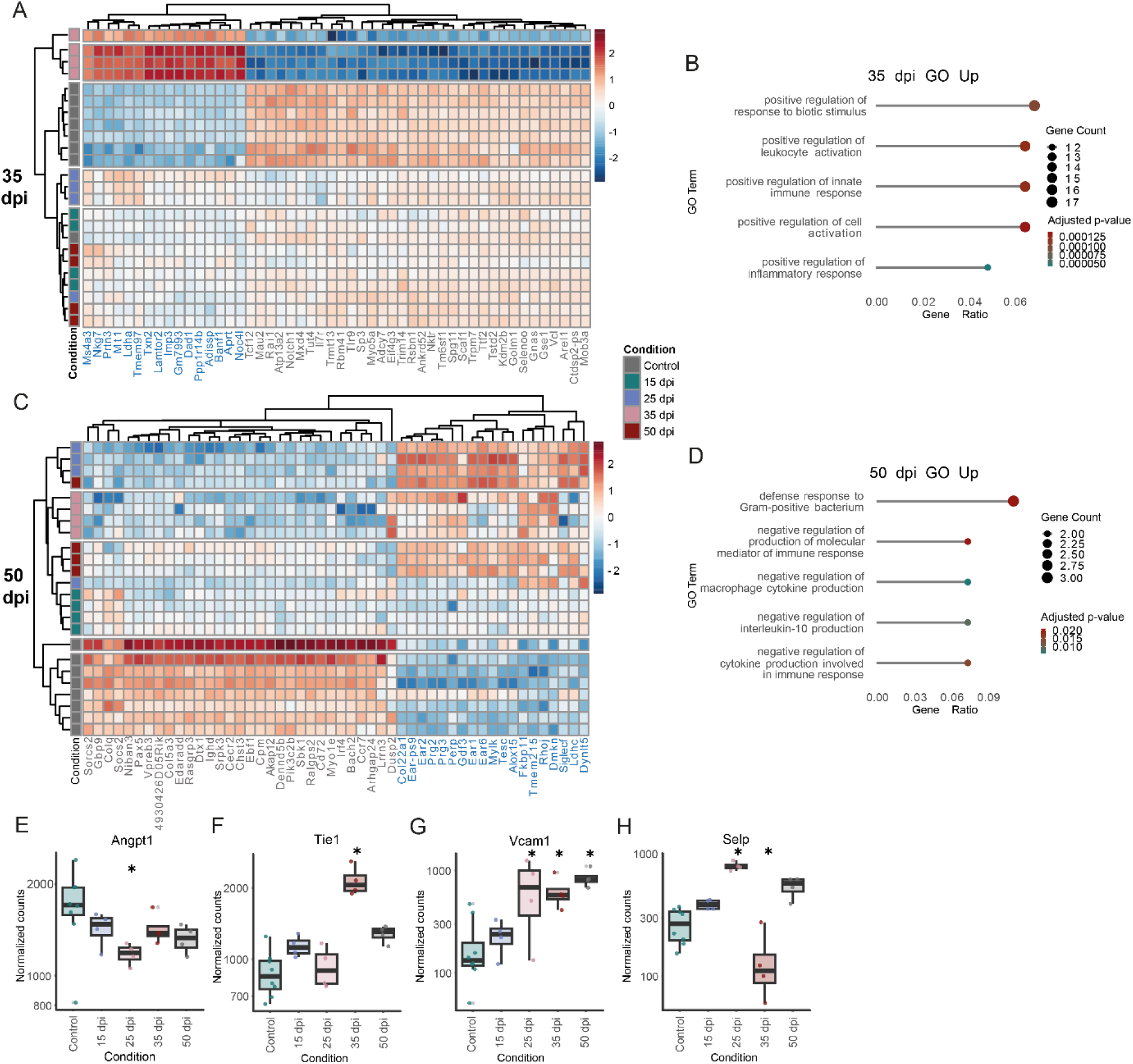
Strong inflammatory gene signature in sorted endothelial cells is induced by CIA in active arthritis. (**A**) Heatmap of DESeq2 normalized gene-expression of top 50 (padj) of 35 dpi DEGs. (**B**) GO analysis of 35 dpi upregulated DEGs (padj < 0.05, log2FC >1.5). (**C**) Heatmap of DESeq2 normalized gene-expression of top 50 (padj) 50 dpi DEGs. (**D**) GO analysis of 50 dpi upregulated DEGs (padj < 0.05, log2FC >1.5). Normalized counts of (**E**) *Angpt1*, (**F**) *Tie1*, (**G**) *Vcam1* and (**H**) *Selp*. Data are presented as boxplots showing the median (line), IQR (box), and whiskers extending to 1.5× IQR. Outliers are shown as individual points. Differential expression analysis was performed using the DESeq2 package, with significance assessed based on p-values adjusted for multiple testing using the Benjamini-Hochberg FDR control method. (n ≥ 4) * padj < 0.05. n ≥ 3.

### Microbiota-specific serum IgG is increased in arthritic mice

In mouse models and individuals with RA, gut microbiota composition was repeatedly shown to differ from healthy controls (39, 40), potentially shaping local immune responses and systemic effects via endothelial cells (41). First, to assess microbiota changes during CIA, stool was collected from naïve controls and CIA mice in the pre-disease, early disease, active disease and remission phase for 16S rRNA-seq. Unweighted UniFrac analysis, sensitive to rare taxa, showed that principal coordinate (PCo) 1 (17.36%) primarily separated at early disease from other samples, and PCo2 (11.92%) separated at pre-disease collected gut microbiota samples (**Fig. 6A**). At the family level, early disease mice exhibited the largest deviations from naïve controls, with decreased *Lachnospiraceae* and increased *Prevotellaceae*, *Rikenellaceae*, and *Muribaculaceae*, while during pre-disease, mice already showed a trend toward lower *Lachnospiraceae* and higher *Lactobacillaceae* (**Fig. 6B**). LEfSe analysis of the pre-, and early disease phases identified taxa with LDA scores >2 showing that at pre-disease, five taxa, including *Tyzzerella* and *Lachnospiraceae*, were enriched; at early disease, five taxa including *Alistipes* and *Rikenellalaceace* were elevated (**Fig. 6C**). To investigate whether these changes reflect systemic bacterial exposure, we measured microbiota-reactive serum IgGs in naïve control mice and CIA mice at pre-disease, early disease and active disease using pooled and allogenic stool coatings. This approach takes advantage of the systemic anti-microbiota IgG repertoire, which can reveal gut bacteria that translocate across intestinal barriers (18). Serum IgG reactivity to gut microbiota was significantly elevated in pre-disease mice and remained significant elevated through disease progression (**Fig. 6D**).

**Fig. 6.**
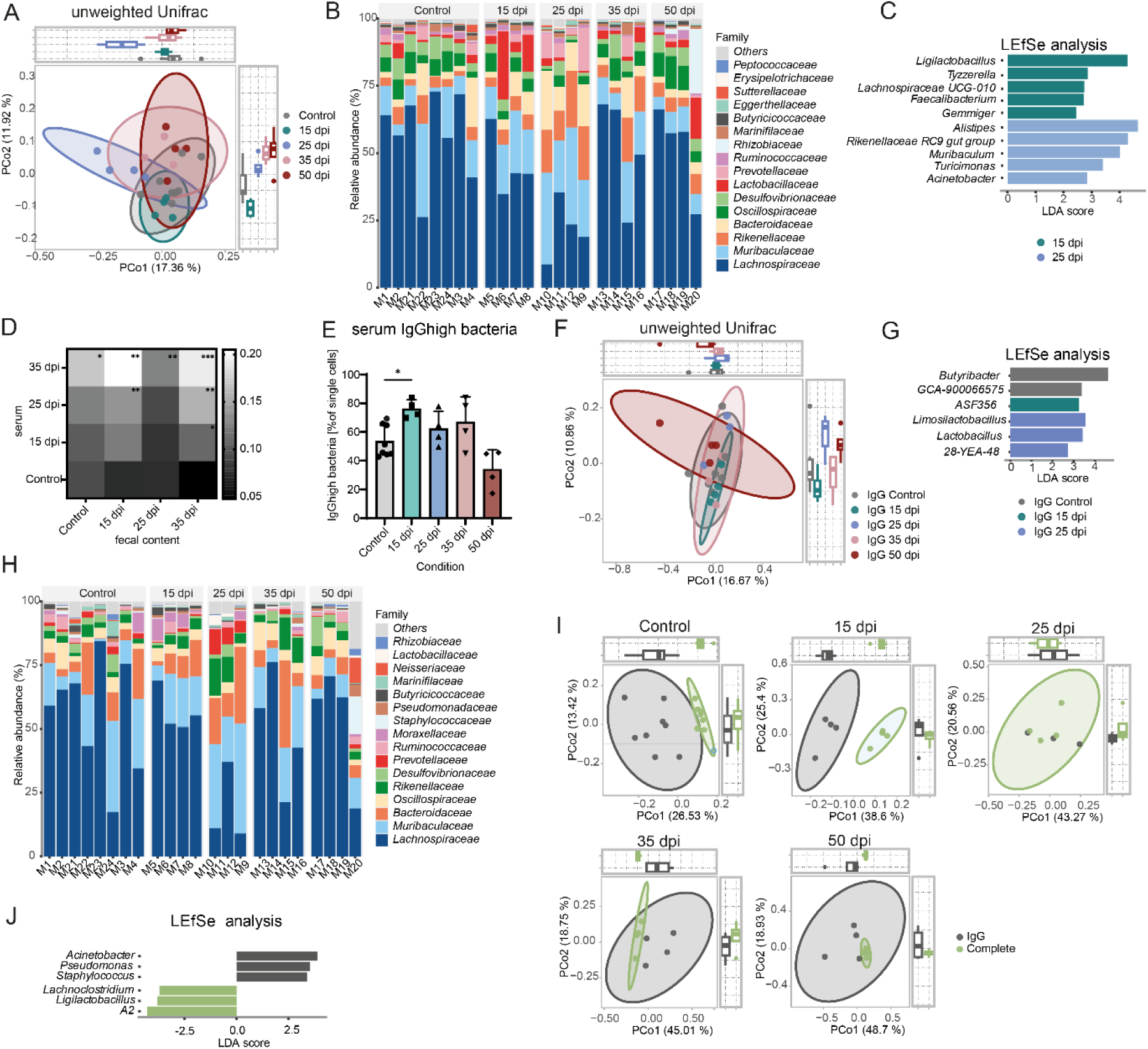
Serum IgG^high^ bacteria are relatively increased in pre-diseased mice and are distinct from the complete bacterial fraction. (**A**) Principal coordinate analysis (PCoA) plot of unweighted Unifrac comparing naïve control mice and different days post immunization (dpi) in CIA. PERMANOVA was used to test for differences between groups with 999 permutations. (**B**) Bar graph showing relative abundance of taxa at family level of the different mice grouped by condition. (**C**) LDA plot of LEfSe analysis from fecal bacteria in naïve control mice and CIA mice at different time points. Features with an LDA score > 2 were considered statistically significant and biologically relevant. (n ≤ 3). (**D**) Levels of Microbiota-reactive IgGs in serum of naïve controls and CIA 15, 25, and 35 dpi measured per ELISA. Values given in arbitrary units (AU). Plates were coated with ileal content from naïve control mice, 15 dpi, 25 dpi and 35 dpi CIA mice. Statistical significance was assessed using two-way repeated measures (RM) ANOVA. (n = 3) * p-value < 0.05 ** p-value < 0.01 *** p-value < 0.001. (**E**) Frequency of serum IgG^high^ bacteria from stool at the different dpi and naïve control mice. (**F**) PCoA plot of unweighted Unifrac comparing naïve control mice and different CIA time points in IgG high fraction. PERMANOVA was used to test for differences between groups with 999 permutations. (**G**) LDA plot of LEfSe analysis from IgG^high^ bacteria in naïve control mice and CIA mice at different time points. Features with an LDA score > 2 were considered statistically significant and biologically relevant. Only the three highest scores from 15 dpi and 25 dpi are shown. (**H**) Bar graph showing relative abundance of taxa at family level of IgG^high^ fraction. (**I**) PCoA plot of unweighted Unifrac comparing IgG positive bacteria and complete bacteria in control and CIA mice at different time points. PERMANOVA was used to test for differences between groups with 999 permutations. (**J**) LDA plot of LEfSe analysis from IgG^high^ bacteria and complete bacteria in naïve control mice. Features with an LDA score > 2 were considered statistically significant and biologically relevant. Only the three highest scores from 15 dpi and 25 dpi are shown. n ≥ 4.

To test if there is a specific gut microbial signature which drives the increase in serum IgG titres, we sorted purified serum IgG^high^ positive bacteria showing significant increases at the pre-disease phase (**Fig. 6E**). Of the sorted serum IgG^high^ bacteria 20.8% were also coated with IgA, which was increased in the early disease phase (Supplementary Fig. 5). When comparing the unweighted Unifrac distance with a focus on rare taxa at different identical timepoint as investigated before there is a significant difference at the pre-disease phase when comparing to control (**Figure 6F**). At genus level *Butyribacter* and *GCA-900066575* define the control group, while *ASF356* is characteristic for the pre-disease state and *Limosilactobacillus*, *Lactobacillus*, as well as *28-YEA-48* define the early disease (**Figure 6G**). On family level, mice at early disease show the highest deviation in the relative abundance levels from the control (**Figure 6H**), with lower levels of *Lachnospiraceae* and a concomitant increase of the families *Muribaculaceae*, *Bacteroidaceae*, *Rikenellaceae* and *Prevotellaceae*. Comparing unweighted UniFrac distances of IgG^high^ vs. total bacteria revealed clear separation in controls and at pre-disease, whereas later time points showed less distinction (**Fig. 6I**, Supplementary Table 5). At steady state, genera contributing most to the difference between IgG^high^ and complete fraction included *Acinetobacter*, *Pseudomonas*, and *Staphylococcus* in the IgG^high^ fraction, and *A2 (Lachnospiraceae)*, *Ligilactobacillus*, and *Lachnoclostridium* in the total fraction (**Fig. 6J**). Together, we show that microbiota-specific serum IgG levels rise already in the pre-disease phase and shift their binding specificity over time.

### Early RA patient gut biopsies show altered predicted cellular interactions similar to arthritic mice

To evaluate the clinical relevance of our findings derived from the CIA model, we further examined ileal gut biopsies from early RA (<1 year disease duration), established RA (>1 year), IBD patients and healthy controls. Patient characteristics including demographic data are shown in Supplementary Fig. 6. Using the established IMC panel (Supplementary Fig. 7, Supplementary Table 6), we detected 525,275 single cells, categorized by E-cadherin (epithelial), Collagen I (stromal), and CD45 (immune) (Supplementary Tables 7 and 8). Rphenograph (k=50) identified 23 clusters, which were assigned to cell types based on marker expression and spatial location (**Figs. 7A and B**, Supplementary Fig. 8). ROIs were pre-classified as epithelial, immune, or mixed epithelial/immune – based on the structural composition of the ROI (**Fig 7B**). Epithelial ROIs showed no major cellular differences across conditions (**Fig. 7C**). Immune and mixed ROIs displayed higher variability between patients (**Figs. 7D and E**), although sample sizes were small. Blinded ROI selection aimed to acquire one epithelial and one immune area per patient. In healthy controls, most ROIs were epithelial (∼77%), while early RA and IBD had relatively more immune ROIs (∼37% and 33%, respectively). In contrast, established RA resembled healthy controls (∼71% epithelial). These findings suggest that while overall cellular composition is similar across conditions, early RA and IBD biopsies may contain more immune-rich regions.

**Fig. 7.**
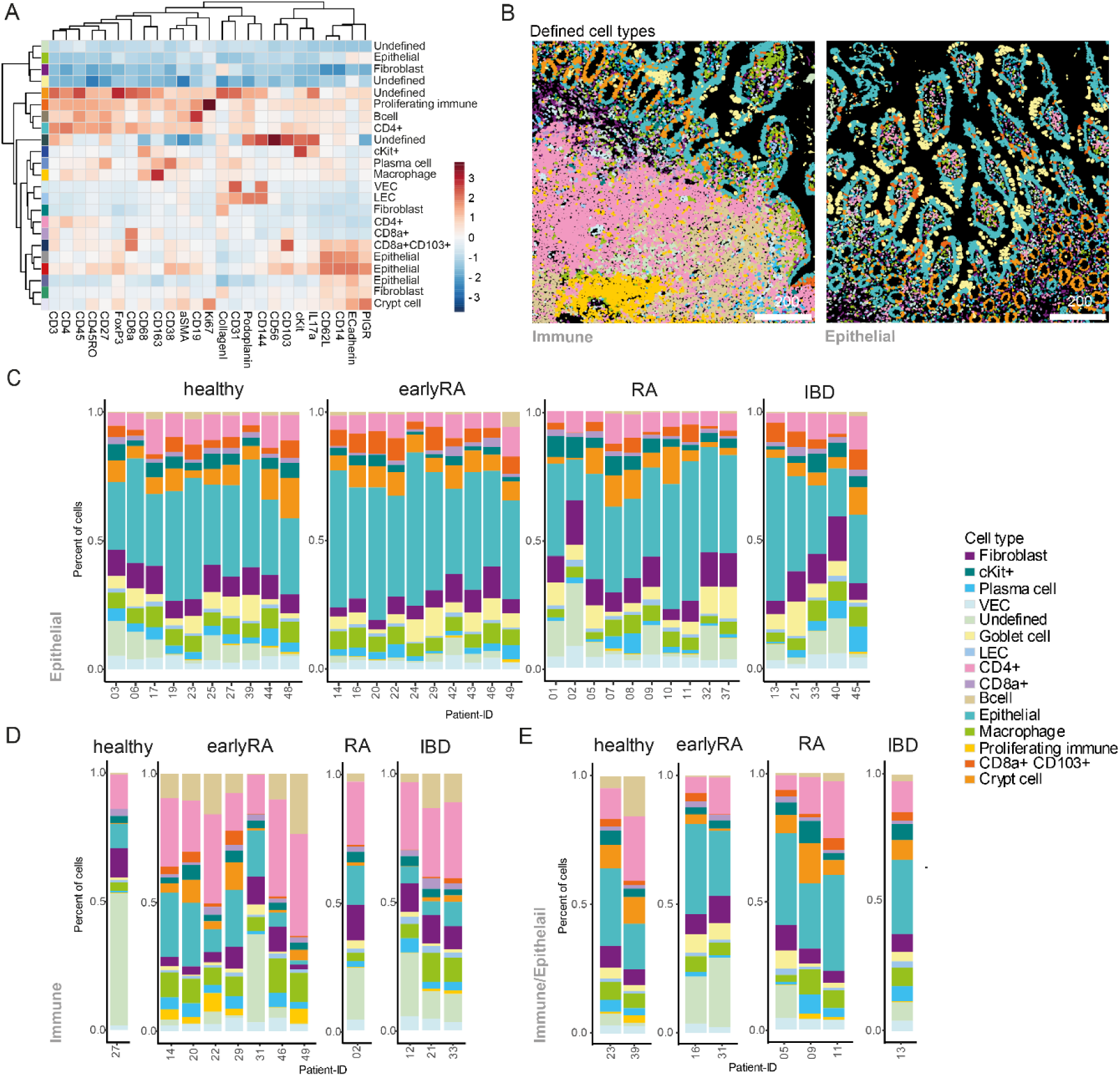
Ileal biopsies from human cohort show no differences in cell type proportions between healthy, early RA, RA and IBD patients. (**A**) Heatmap of normalized marker expression with color codes for 23 phenograph clusters and the respective defined cell types. The heatmap colors represent the z-score of the average expression of a given marker for each identified cluster. (**B**) Cell plots from two example ROIs of the 15 based on marker expression and localization defined cell types. Bar graphs showing the mean percentage of cells in the different conditions belonging to the different defined cell types per patient ID in samples of tissue type. (**C**) Epithelial, (**D**) Immune and (**E**) Immune/Epithelial.

We used Delaunay triangulation (max_dist = 20) to predict cell–cell interactions between cell types (**Fig. 8A**). Based on these predicted interactions, cells were assigned to six CNs, whose relative proportions remained stable throughout the disease course (**Figs. 8B and C**). Epithelial tissue samples were enriched in CN2, immune tissues in CN3 and CN4, and mixed tissues showed intermediate distributions (**Fig. 8C**). CN1 mainly contained crypt cells, CN2 epithelial cells, CN3 CD4^+^ T cells, CN4 undefined cells, CN5 a mix of fibroblasts, macrophages and VECs, and CN6 epithelial cells with CD8a^+^CD103^+^ cells (**Fig. 8D**). Looking at the mean cell type proportions in the different CNs, there is a high homogeneity between the tissue types (**Fig. 8D**). In CN1, the immune-type, the healthy condition is represented by a single sample, showing a high proportion of undefined (cell type could not be characterized based on marker expression and spatial location), epithelial, and goblet cells. Cell–cell interactions were analysed in the early RA group, corresponding to the pre- and early CIA stage, which showed the greatest intestinal changes. Compared to healthy controls, CN1 showed reduced fibroblast–cKit⁺ predicted interactions, while CN2 exhibited increased CD8a^+^–CD4^+^ predicted interactions. In CN3, interactions were predicted to be decreased between plasma–undefined cells, CD4^+^–cKit⁺ cells, and CD8a^+^–cKit⁺ cells. CN4 showed increased predicted interactions of B cells with CD4^+^, proliferating immune, and undefined cells. The largest changes occurred in CN5, with reduced CD4–cKit⁺, CD8a^+^–cKit⁺, and fibroblast–cKit⁺ predicted interactions, and increased CD8a^+^CD103^+^–macrophage, macrophage–macrophage, and macrophage–proliferating immune predicted interactions. In CN6, CD8a^+^CD103^+^ cells had elevated predicted interactions with CD8a^+^ cells (summarized in **Fig. 8E**).

**Fig. 8.**
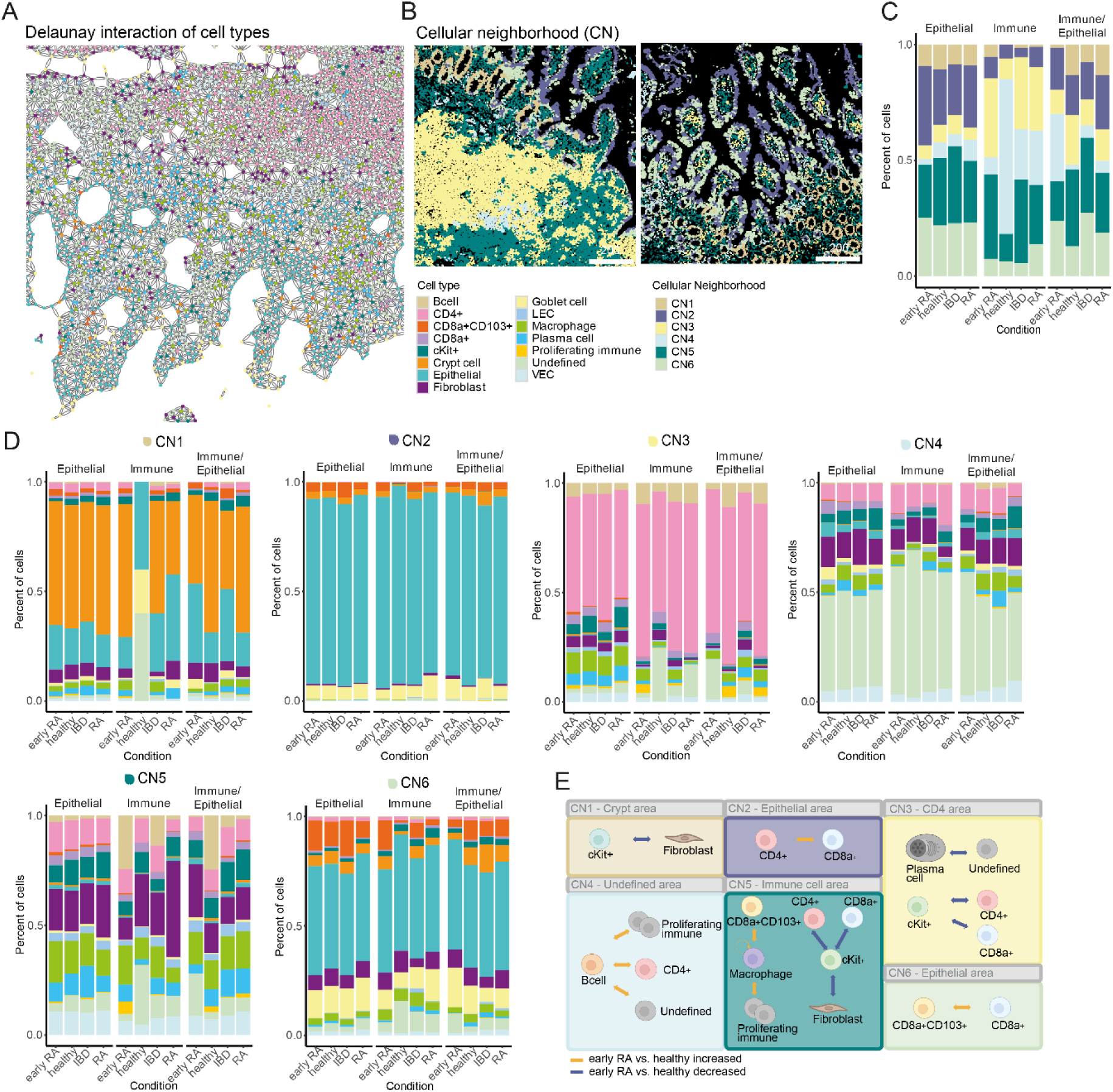
Human early RA patients show changes in the predicted interaction of immune cells in cellular compartments. (**A**) Delaunay triangulation-based interaction prediction of the different cell types in one example ROI. (**B**) Cell plots of example ROIs of cellular neighborhoods (CN)1 to CN6 detected in the human data-set. (**C**) Bar graph showing the mean percentage of cells of the different CNs in the different conditions. (**D**) Bar graphs showing the percentage of cells of the different cell types in CN1 to CN6 (upper left to lower right) in the different conditions grouped by tissue type. (**E**) Graphical illustration of significantly changed predicted cellular interactions (one-way ANOVA p<0.05 & Tukey HSD p<0.05) in the different CNs comparing early RA patients with healthy controls. Predicted interactions (based on Delaunay triangulation) of the different cell types in one CN per ROI grouped by condition Number of Interactions between cell types in one CN per ROI were normalized on the cell number in the CN per ROI (nCNinteractions/nCNcellcount *100). Orange arrows indicates increased predicted interaction of the cell types. Blue arrows indicate reduced predicted interaction of the cell types.

### Early RA patients show signs of lymphocyte activation in the gut similar to IBD

IMC data revealed no major differences in cell type composition across conditions, but predicted interactions between cell types within CNs varied. To further explore cellular states, bulk RNA-seq. was performed on ileal biopsies from the same cohort. PCA showed that sample variance was largely independent of condition or sex, with PC1 explaining 56% of variance (**Fig. 9A**). The top 200 genes influencing PC1 were associated with cellular components such as the brush border and apical cell regions, indicating structural differences between samples (**Fig. 9B**). Expression of *CDH1* (E-Cadherin) and *PTPRC* (CD45) suggested patient-specific tissue-type differences, consistent with IMC observations (**Fig. 9C**). Comparing normalized gene expression to healthy controls, the largest differences were observed in early RA (99 DEGs: 90 up, 9 down), with upregulated genes involved in lymphocyte differentiation and B cell activation (**Figs. 9D, 9F and 9G**). IBD samples showed 72 DEGs (61 up, 11 down), associated with T cell activation and chemokine-mediated signalling (**Figs. 9E and 9H**). Looking for similar patterns in the regulated genes in the groups, most overlap is observed in the early RA and IBD group with 16 common genes (**Fig. 9F**). Among these, 14 are upregulated in both groups and include the genes *LTB*, *CCR7*, *CCR4*, CCL22, C3 and *CTLA4*, of which some are involved in the response to chemokines and positive regulation of inflammatory response to antigenic stimulus (**Fig. 9I**).

**Fig. 9.**
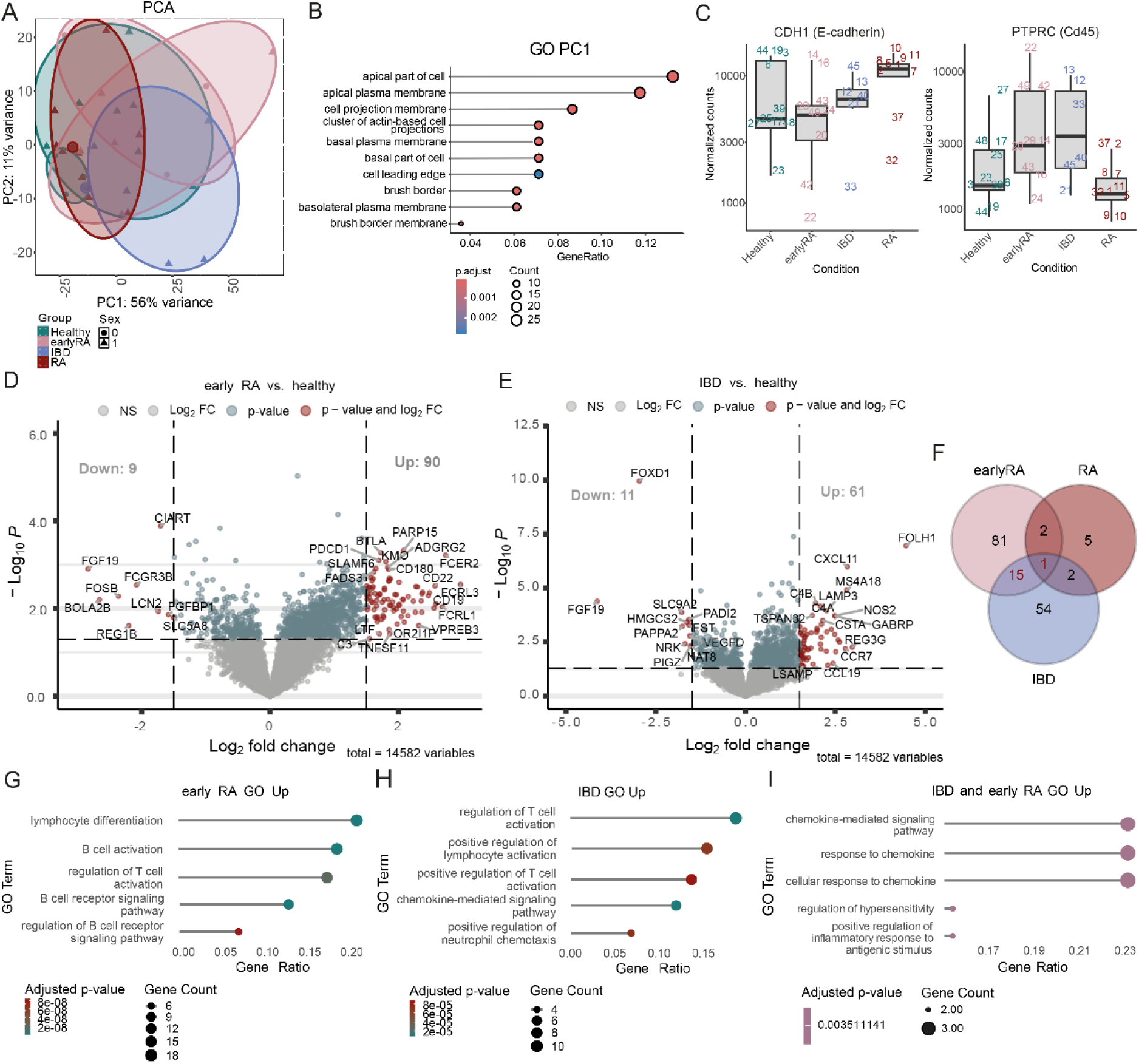
Human early RA patients how signs of lymphocyte activation in the gut. (**A**) Principal component analysis (PCA) plot of top 500 variable genes with sex and condition as grouping variables. (**B**) GO enrichment analysis (only cellular component, maxGSSize = 500) of top 200 genes of PC1. Padj was calculated using the Benjamini Hochberg method. (**C**) Normalized counts (DESeq2) of CDH1 and PTPRC comparing the different conditions (Healthy, early RA, IBD, RA). Numbers refer to different patient IDs. Volcano plots of Log2 fold change (FC) values of mRNA sequencing analysis from ileal biopsies from human cohort including (**D**) early RA and (**E**) IBD compared to healthy subjects. Gates were set at a p-value < 0.05 and a log2 FC > 1.5. Genes marked in red display genes above log2 FC gate and p-value gate. Blue dots represent genes above p-value gate and below the log2 FC gate. Grey dots display genes below p-value gate. (**F**) Venn diagram of regulated genes (above p-value gate and log2 FC gate). Results of GO term analysis of upregulated genes in (**G**) early RA group and (**H**) IBD group compared to healthy subjects. (**I**) results of GO term analysis of the genes upregulated both in early RA group and IBD group compared to healthy subjects. Differential expression analysis was performed using the DESeq2 package, with significance assessed based on p-values adjusted for multiple testing using the Benjamini-Hochberg FDR control method.

## Discussion

Using complementary mouse and human datasets, we demonstrate that intestinal endothelial and immune alterations precede or coincide with the earliest phases of inflammatory arthritis. These findings support a model in which mucosal barrier dysfunction actively participates in disease initiation rather than representing a secondary consequence of systemic inflammation. In the CIA model, *in vivo* imaging revealed increased endothelial permeability in the intestine during the preclinical phase, preceding detectable synovial inflammation. Together with the published results on the epithelial leakiness in arthritis (5–7), such early vascular leakage may further facilitate the paracellular passage of microbial molecules systemically and permit immune cell infiltration into the lamina propria.

Endothelial leakiness is well documented in IBD (33) and in synovial vasculature during RA, where cytokines and growth factors disrupt junctional integrity and promote leukocyte influx (42). Here, we show that intestinal endothelial cells undergo the most profound transcriptional remodelling already in the pre-disease phase and not during the active synovial inflammation. Upregulated genes associated with endothelial activation and leukocyte trafficking (e.g., *Sele, Madcam1, Glycam1, Ackr1*) suggest a primed vascular state that may shape subsequent mucosal immune activity.

Spatial neighbourhood analysis of the IMC data from the intestine revealed a notable increase in predicted interactions among epithelial cells, vascular endothelial cells, IELs and macrophages. Consistently, in early RA patient biopsies, CD8^+^CD103^+^ IEL-like cells showed enhanced predicted interactions with macrophages and CD8^+^ T cells, highlighting similar mucosal immune signatures in RA and CIA.

In contrast to the intestine, bone marrow endothelial cells exhibited a strong type IFN-I signature in the pre-disease phase, including upregulation of *Ifi44*, which was also shown to be elevated in synovial tissues and peripheral blood of RA patients (43, 44). IFN-I signatures are consistently enriched in early RA, even before arthritis onset (45), and are linked to bacterial translocation through their complex roles in the immune response to bacterial infections (46) IFN-I can increase endothelial permeability and downregulate *Angpt1*, a stabilizing angiopoietin (47). Concordantly, CIA bone endothelium showed reduced *Angpt1* and increased *Vcam1* and *Selp* expression, markers observed in IBD for endothelial dysfunction (48, 49), indicating endothelial activation and enhanced leukocyte trafficking. Although limited IFN-I-related genes were upregulated in the intestine, transcriptional remodelling was substantially more pronounced in the bone marrow tissue during active disease. While intestinal endothelial cells returned to baseline transcriptional profiles during remission, bone marrow endothelial cells retained distinct signatures, implying tissue-specific resolution dynamics.

In human intestinal biopsies, immune-cell–enriched regions were more frequent in early RA than in established RA mimicking findings in IBD patients, showing increased CD4 – B cell predicted interactions and pointing to early mucosal lymphoid activation. The increased predicted interactions between CD8a^+^CD103^+^ cells and the higher frequency of macrophage contacts in the immune cell area could indicate early formation or reorganization of lymphoid structures in the gut, such as isolated lymphoid follicles or tertiary lymphoid structures, which play a key role in inflammation and autoimmunity (50).

Subsequent bulk RNA sequencing of identical intestinal biopsies showed significant enrichment of genes associated with response to chemokines and positive regulation of inflammatory response to an antigenic stimulus in early RA, including *LTB*, *CCR7*, *CCR4*, and *CTLA4* - genes that also elevated in IBD (51–54) and linked to the formation of isolated lymphoid follicles or tertiary lymphoid structures (55).

In contrast to RA, IBD is well recognized to involve gut-barrier disruption and heightened immune activation, processes strongly shaped by the intestinal microbiota. Levels of faecal bacteria coated with endogenous IgG are increased in both CD and UC (56, 57) and correlate with disease activity (58, 59). Prior studies show that systemic IgG responses to selected commensals rise following bacterial translocation and that IgG-repertoire profiling can identify translocating taxa. Barrier impairment also drives divergence between systemic IgG and mucosal IgA repertoires, with IgG becoming enriched for translocating and proinflammatory organisms (18). Consistent with this, we observed a progressive increase in serum IgG responses to microbial antigens over the course of CIA, with IgGs recognizing intact bacteria peaking in the pre-disease phase. These antibodies likely target conserved surface structures such as β-glucans or lipopolysaccharides, indicative of T cell–independent class switching driven by pattern-recognition receptor signalling (60, 61) . The subsequent rise during the disease phase in IgGs reactive to bacterial fragments suggests the onset of T cell–dependent responses, potentially reflecting adaptive immunity to invasive or adherent bacterial populations (60, 62).

Longitudinal binding assays and 16S rRNA sequencing of sorted IgG bound bacteria revealed an expansion of the IgG^high^ bacterial fraction and a convergence between rare-taxa composition and the overall microbial community, indicating that previously low-abundance taxa become increasingly targeted as immune activation intensifies and barrier function deteriorates. Given that rare taxa are known to exert disproportionate immunostimulatory effects (63), their enhanced recognition may contribute to early immune adaptation and shape the trajectory of autoimmune inflammation.

In early phases of arthritis, serum IgA coating bacteria were highest and coincided with an increased abundance of adhesive Lactobacillus species. In the later disease phase IgA and IgG responses to overlapping taxa are characteristic of polyreactive mucosal plasma cells, which can be generated independently of dietary or microbial antigens (64).

Together, these findings support a model in which early endothelial activation compromises barrier integrity, enabling the translocation of bacterial products and the induction of both polyreactive and antigen-specific IgG responses. These antibodies may, in turn, amplify systemic inflammation through molecular mimicry or immune-complex formation, thereby contributing to arthritis initiation.

## Methods

### Sex as a biological variable

The CIA experiments were examined exclusively in female mice because RA is more common in females (65). The effect of microbiota reactive IgGs on the differentiation of osteoclasts was examined in male and female animals, and similar findings are reported for both sexes. Our human study examined male and female patients. These were analysed as pooled data of both sexes without assessment of sex-specific differences.

### Mice

All mice were maintained under specific pathogen-free conditions at the local animal facility or the at the Präklinisches Experimentelles Tierzentrum (PETZ), Erlangen, Germany and approved by the local ethics authorities of the Regierung of Unterfranken (00063105-2-4; 55.2-2532.2-630; TS-7/2021). Five to six-week-old wildtype female DBA/1J were purchased from Janvier (Janvier Labs, Le Genest-Saint-Isle, France) and acclimated for one week, followed by two weeks of co-housing before starting the experiment. The animals received water and standard chow (Sniff Spezialdiäten GmbH, Soest, Germany) ad libitum.

### Collagen-induced arthritis (CIA)

CIA was induced in 8-week-old female DBA/1J mice by subcutaneous injection at the base of the tail with 100 µl of 0.25 mg chicken type II collagen (CII; Chondrex) in Complete Freund’s Adjuvant (CFA; Difco Laboratory) containing 5 mg/ml killed Mycobacterium tuberculosis (H37Ra). Mice were re-challenged after 21 days intradermal immunization in the base of the tail with 100 µl of 0.25 mg chicken type II collagen (CII; Chondrex) in incomplete Freund adjuvant. The paws were evaluated for joint swelling three times per week. Each paw was individually measured for paw thickness using a caliper. All CIA mice developed clinical symptoms. No exclusions due to deviating clinical scores were performed.

### Treatment with α4β7 antibody and Imatinib-mesylate

Vedolizumab (α4β7 antibody) was administered by intraperitoneal injections (5 mg/kg bodyweight) 3 times per week starting 0 dpi until 21 dpi of CIA. Imatinib-mesylate (Merck, Darmstadt, Germany) was solved in DMSO (100 mg/ml) and diluted in PBS for daily gavage from 15 dpi until 25 dpi (50 mg/kg bodyweight).

### Human study cohort

Intestinal biopsies from 40 individuals within the research project Intestinal link to Rheumatoid Arthritis (IntestRA) were acquired. Ethical approval was granted by the Swedish Ethical Review Authority on 13^th^ January 2016 (number 2015-415). An amendment for the permission to send samples to Germany for a collaborative project has been approved on April 12th 2021 (number 2021-01477). Written informed consents have been acquired from all participants. Biopsies from 37 persons were sent and analyzed within the project A1 of CRC369.

### *In vivo* imaging of colonic endothelial leakiness

In vivo assessment of colonic vascular permeability was performed as described previously (33, 66) in collaboration with AG Stürzl at the Translational Research Center (TRC, Erlangen, Germany) as described in **Supplementary Methods**.

### Imaging mass cytometry (IMC)

The protocol for the IMC workflow including tissue preparation and data analysis for mouse and human tissue are provided in the **Supplementary Methods**. The used antibodies are listed in **Supplementary Table 1** for mouse and **Supplementary Table 6** for human tissue.

### Bulk RNA sequencing analysis of murine intestinal endothelial cells

Intestinal endothelial cells were isolated as previously described by Tisch et al. (2022) with minor modifications. The detailed protocol for the isolation of intestinal endothelial cells, RNA extraction and bulk RNA Seq.-analysis are provided in the supplementary methods.

### Bulk RNA sequencing analysis of murine bone endothelial cells

The protocol for the isolation of bone endothelial cells, RNA extraction and bulk RNA Seq-analysis are provided in the **Supplementary Methods.**

### 16S rRNA analysis of serum IgG bound fecal bacteria

The protocol for stool preparation, staining, FACS and 16S rRNA analysis are provided in the Supplementary Methods.

### Quantification of commensal bacteria-reactive IgG using ELISA

The protocol for stool preparation and ELISA is provided in the **Supplementary Methods.**

### Bulk RNA sequencing of human ileal biopsies

The protocol for tissue preparation, RNA extraction and bulk mRNA sequencing are provided in the **Supplementary Methods**.

### Data analysis and statistics

Statistical analyses were performed using Prism 9 software (GraphPad) and RStudio (R version 4.3.3). For comparisons between two independent groups with not normally distributed data, a Mann-Whitney U test was performed. For Comparisons between two dependent groups with not normally distributed data, a Wilcoxon matched-pairs signed-rank test was performed. For comparisons between two groups with normally distributed data unpaired or paired, two-tailed, Student’s t test was performed. Comparisons between more than two groups were performed using one-way ANOVA and post-hoc Tukey’s or Dunnett’s multiple comparison test. For the mRNA sequencing-analysis the DESeq2 package was used, which controls for False Discovery Rate (FDR) by using the Benjamini-Hochberg (BH) procedure with the FDR set to 0.05. For the GO enrichment, FDR control was performed using the BH procedure. Details on the statistical analysis are listed in the figure legends and the data analysis part in the IMC, 16S rRNA and bulk mRNA sequencing analysis. * p < 0.05; ** p < 0.01; *** p < 0.001; **** p < 0.0001

## Data Availability

All relevant data are available from the authors upon reasonable request. The source data underlying Figures 1-9 and Supplementary Figures 1-8 are provided as source data file.

## Author contributions

E.S., N.O, G.S., B.W. and M.M.Z. designed the project, interpreted results and wrote the manuscript. E.S., N.O, S.A. and H.D. performed most of the work, analysed data, edited the manuscript and made figure panels. E.S., N.O, S.A., H.D., M.F., E.N. and L.S. acquired data and provided help with multiple experiments. G.F. provided the microbiota specific IgG sequencing setup and platform. A.K. and A.S. provided human gut biopsy samples. M.M.B. supported the IMC analysis. E.S. and K.S. wrote the animal licence approvals. M.M.Z supervised the project and provided funding. All the authors revised and approved the manuscript.

## Funding support

The research presented in this study was supported by the Deutsche Forschungsgemeinschaft (DFG, German Research Foundation) through funding provided for project B04 within the Research Training Group “Immunomicrotope” (GRK 2740/447268119), as well as via DFG funding for DFG - SFB/TRR369 DIONE (501752319), grant 543936725 ZA 899/22-1, and DFG - TRR 417 (project ID 540805631; subprojects P06 and S01 to EN). Additional support was received from the IZKF project P144. AK acknowledges funding from the King Gustaf V’s 80-year foundation, the Swedish Research Council, the Swedish Rheumatism Association, and ALF grants from region Östergötland.

## Acknowledgements

We thank Dr. Stefan Wirtz and the MACE facility at Medical Clinic 1, University Hospital Erlangen, Germany, for 16S rRNA sequencing. We also thank Dr. Aleix Rigau for valuable support with the IMC at Medical Clinic 3, University Hospital Erlangen, Germany. We are grateful to all members of the laboratories at Medical Clinic 3, University Hospital Erlangen, Germany, for their support and insightful discussions. We thank the PharmBio EV facility at FAU Erlangen (Lorenzo Sana, Leila Pourtalebijahromi and Gregor Fuhrmann) for the use of their flow cytometer (DFG grant No. 511550821). We also thank Dr Daniel Sjöberg at Falun Hospital, Sweden, for performing colonoscopies and obtaining human intestinal biopsies.

## Competing interest

The authors declare no competing interests.

## Supplementary Material

**Supplementary Figs. 1 - 8**

**Supplementary Tables 1 – 9**

## Supplementary Methods

## Notes

The authors have declared that no conflict of interest exists.

### Competing Interest Statement

The authors have declared no competing interest.

