## Supplementary Tables for "Microbiota-specific serum IgG links gut and joints through immune–endothelial crosstalk in arthritis"

**Table 1. Imaging Mass Cytometry panel – mouse.**

| Mass | Metal | Target | Clone | Company | DF |
|---|---|---|---|---|---|
| 89 | Y | CD45 | 30-F11 | Fluidigm | 400 |
| 141 | Pr | Alpha SMA | 1A4 | Fluidigm | 500 |
| 142 | Nd | CD11c | N418 | Fluidigm | 300 |
| 143 | Nd | Vimentin | D21H3 | Fluidigm | 1000 |
| 144 | Nd | CD103 | QA17A24 | Biolegend | 200 |
| 145 | Nd | CD4 | RM4-5 | Fluidigm | 400 |
| 146 | Nd | CD5 | 537.3 | Fluidigm | 500 |
| 147 | Sm | CD206 | C068C2 | Biolegend | 200 |
| 148 | Nd | CD11b/Mac-1 | M1/70 | Fluidigm | 200 |
| 149 | Sm | CD19 | 6D5 | Fluidigm | 200 |
| 150 | Nd | CD27 | LG.3A10 | Fluidigm | 200 |
| 151 | Eu | CD25 | 3C7 | Fluidigm | 200 |
| 152 | Sm | VEGFR1 | polyclonal | R&D systems | 200 |
| 153 | Eu | CD8a | 536.7 | Fluidigm | 200 |
| 154 | Sm | CD163 | S15049I | Biolegend | 100 |
| 155 | Gd | VEGFR2 | Avas12 | Biolegend | 200 |
| 156 | Gd | Madcam-1 | MECA-367 | Biolegend | 200 |
| 158 | Gd | Foxp3 | FJK-16S | Fluidigm | 100 |
| 159 | Tb | F4/80 | BM8 | Fluidigm | 400 |
| 160 | Gd | Lyve-1 | 223322 | R&D systems | 200 |
| 161 | Dy | CD68 | FA-11 | Biolegend | 200 |
| 162 | Dy | T-bet | 4B10 | Biolegend | 300 |
| 163 | Dy | CD54 (ICAM-1) | YN1/1.7.4 | Fluidigm | 300 |
| 164 | Dy | CD69 | H1.2F3 | Biolegend | 200 |
| 165 | Ho | CD31 (PECAM-1) | 390 | Fluidigm | 300 |
| 166 | Er | C-kit (Cd117) | 2B8 | Fluidigm | 100 |
| 167 | Er | VE-Cadherin | BV13 | Biolegend | 100 |
| 168 | Er | Ki-67 | B56 | Fluidigm | 300 |
| 169 | Tm | Collagen | poly | Fluidigm | 500 |
| 170 | Er | PV-1 | MECA-32 | Abcam | 200 |
| 171 | Yb | CD3 | 145-2C11 | Fluidigm | 200 |
| 172 | Yb | GFAP | polyclonal | Abcam | 800 |
| 173 | Yb | RORgammaT | W19344C | Biolegend | 100 |
| 174 | Yb | Ly6G | 1A8 | Biolegend | 200 |
| 175 | Lu | CD127/IL-7Ra | A7R34 | Fluidigm | 200 |
| 176 | Yb | Desmin | Y66 | Abcam | 1000 |
| 195 | Pt | ICSK1 | | Fluidigm | 400 |
| 196 | Pt | ICSK2 | | Fluidigm | 400 |
| 198 | Pt | ICSK3 | | Fluidigm | 400 |
| 209 | Bi | MHCII (I-A/I-E) | M5/114.15.2 | Fluidigm | 500 |

**Table 2. Samples list with cell numbers of via IMC analysed ileal tissue – mouse.**

| Condition | Mouse | Sample | ROI | ROI size | Cells per ROI | Total cells per Mouse | Total cells per Group |
|---|---|---|---|---|---|---|---|
| Control | 1 | 1 | ES-33 day0 m1_002 | 1000x1100 | 10375 | 28537 | 87297 |
| Control | 1 | 2 | ES-33 day0 m1_003 | 1000x1100 | 9965 | | |
| Control | 1 | 3 | ES-33 day0 m1_005 | 1000x1100 | 8197 | | |
| Control | 2 | 4 | ES-33 day0 m2_001 | 1000x1100 | 9594 | 29136 | |
| Control | 2 | 5 | ES-33 day0 m2_002 | 1100x1000 | 9673 | | |
| Control | 2 | 6 | ES-33 day0 m2_003 | 1000x1100 | 9869 | | |
| Control | 3 | 7 | ES-33 day0 m3_001 | 1000x1100 | 10662 | 29624 | |
| Control | 3 | 8 | ES-33 day0 m3_002 | 1000x1100 | 9039 | | |
| Control | 3 | 9 | ES-33 day0 m3_003 | 1100x1000 | 9923 | | |
| 5 dpi | 4 | 10 | ES-33 day5 m1_001 | 1100x1000 | 9908 | 28327 | 87453 |
| 5 dpi | 4 | 11 | ES-33 day5 m1_002 | 1100x1000 | 9678 | | |
| 5 dpi | 4 | 12 | ES-33 day5 m1_003 | 1100x1000 | 8741 | | |
| 5 dpi | 5 | 13 | ES-33 day5 m2_001 | 1100x1000 | 10615 | 29950 | |
| 5 dpi | 5 | 14 | ES-33 day5 m2_002 | 1000x1100 | 9842 | | |
| 5 dpi | 5 | 15 | ES-33 day5 m2_003 | 1100x1000 | 9493 | | |
| 5 dpi | 6 | 16 | ES-33 day5 m3_001 | 1100x1000 | 10729 | 29176 | |
| 5 dpi | 6 | 17 | ES-33 day5 m3_002 | 1000x1100 | 8603 | | |
| 5 dpi | 6 | 18 | ES-33 day5 m3_003 | 1100x1000 | 9844 | | |
| 10 dpi | 7 | 19 | ES-33 day10 m1_001 | 1100x1000 | 8771 | 24999 | 80752 |
| 10 dpi | 7 | 20 | ES-33 day10 m1_002 | 1000x1100 | 8735 | | |
| 10 dpi | 7 | 21 | ES-33 day10 m1_003 | 1100x1000 | 7493 | | |
| 10 dpi | 8 | 22 | ES-33 day10 m2_001 | 1000x1100 | 9791 | 28284 | |
| 10 dpi | 8 | 23 | ES-33 day10 m2_002 | 1100x1000 | 8886 | | |
| 10 dpi | 8 | 24 | ES-33 day10 m2_003 | 1000x1100 | 9607 | | |
| 10 dpi | 9 | 25 | ES-33 day10 m3_001 | 1000x1100 | 8704 | 27469 | |
| 10 dpi | 9 | 26 | ES-33 day10 m3_002 | 1000x1100 | 9508 | | |
| 10 dpi | 9 | 27 | ES-33 day10 m3_003 | 1000x1100 | 9257 | | |
| 15 dpi | 10 | 28 | ES-33 day15 m1_001 | 1000x1100 | 9041 | 23389 | 72554 |

| | | | | | | | |
|---|---|---|---|---|---|---|---|
| 15 dpi | 10 | 29 | ES-33 day15 m1_002 | 1100x1000 | 8652 | | |
| 15 dpi | 10 | 30 | ES-33 day15 m1_003 | 1000x1100 | 5696 | | |
| 15 dpi | 11 | 31 | ES-33 day15 m2_001 | 1100x1000 | 7335 | | |
| 15 dpi | 11 | 32 | ES-33 day15 m2_2_001 | 1000x1100 | 12467 | 26819 | |
| 15 dpi | 11 | 33 | ES-33 day15 m2_2_002 | 1100x1000 | 7017 | | |
| 15 dpi | 12 | 34 | ES-33 day15 m3_001 | 1000x1100 | 5878 | | |
| 15 dpi | 12 | 35 | ES-33 day15 m3_002 | 1000x1100 | 7838 | 22346 | |
| 15 dpi | 12 | 36 | ES-33 day15 m3_003 | 1000x1100 | 8630 | | |
| 20 dpi | 13 | 37 | ES-33 day20 m1_001 | 1000x1100 | 9856 | | |
| 20 dpi | 13 | 38 | ES-33 day20 m1_005 | 1100x1000 | 8253 | 24765 | |
| 20 dpi | 13 | 39 | ES-33 day20 m1_007 | 1000x1100 | 6656 | | |
| 20 dpi | 14 | 40 | ES-33 day20 m2_001 | 1100x1000 | 12868 | | |
| 20 dpi | 14 | 41 | ES-33 day20 m2_002 | 1000x1100 | 13715 | 39608 | 87037 |
| 20 dpi | 14 | 42 | ES-33 day20 m2_003 | 1100x1000 | 13025 | | |
| 20 dpi | 15 | 43 | ES-33 day20 m3_001 | 1000x1100 | 7858 | | |
| 20 dpi | 15 | 44 | ES-33 day20 m3_002 | 1000x1100 | 8800 | 22664 | |
| 20 dpi | 15 | 45 | ES-33 day20 m3_003 | 1000x1100 | 6006 | | |
| 25 dpi | 16 | 46 | ES-33 day25 m1_001 | 1000x1100 | 9620 | | |
| 25 dpi | 16 | 47 | ES-33 day25 m1_002 | 1100x1000 | 11042 | 32284 | |
| 25 dpi | 16 | 48 | ES-33 day25 m1_003 | 1000x1100 | 11622 | | |
| 25 dpi | 17 | 49 | ES-33 day25 m2_001 | 1000x1100 | 12657 | | |
| 25 dpi | 17 | 50 | ES-33 day25 m2_002 | 1000x1100 | 13361 | 36493 | 97527 |
| 25 dpi | 17 | 51 | ES-33 day25 m2_003 | 1100x1000 | 10475 | | |
| 25 dpi | 18 | 52 | ES-33 day25 m3_001 | 1000x1100 | 7828 | | |
| 25 dpi | 18 | 53 | ES-33 day25 m3_002 | 1100x1000 | 10691 | 28750 | |
| 25 dpi | 18 | 54 | ES-33 day25 m3_003 | 1000x1100 | 10231 | | |
| 30 dpi | 19 | 55 | ES-33 day30 m1_001 | 1000x1100 | 9315 | | |
| 30 dpi | 19 | 56 | ES-33 day30 m1_002 | 1100x1000 | 10008 | 32625 | |
| 30 dpi | 19 | 57 | ES-33 day30 m1_003 | 1000x1100 | 13302 | | |
| 30 dpi | 20 | 58 | ES-33 day30 m2_001 | 1000x1100 | 7158 | | 84545 |
| 30 dpi | 20 | 59 | ES-33 day30 m2_002 | 1000x1100 | 10954 | 27959 | |
| 30 dpi | 20 | 60 | ES-33 day30 m2_003 | 1000x1100 | 9847 | | |

| | | | | | | | |
|---|---|---|---|---|---|---|---|
| 30 dpi | 21 | 61 | ES-33 day30 m3_001 | 1000x1100 | 10464 | | |
| 30 dpi | 21 | 62 | ES-33 day30 m3_002 | 1100x1000 | 6022 | 23961 | |
| 30 dpi | 21 | 63 | ES-33 day30 m3_003 | 1000x1100 | 7475 | | |
| 35 dpi | 22 | 64 | ES-33 day35 m1_001 | 1000x1100 | 11938 | | |
| 35 dpi | 22 | 65 | ES-33 day35 m1_002 | 1100x1000 | 11536 | 34705 | |
| 35 dpi | 22 | 66 | ES-33 day35 m1_003 | 1000x1100 | 11231 | | |
| 35 dpi | 23 | 67 | ES-33 day35 m2_003 | 1100x1000 | 11934 | | |
| 35 dpi | 23 | 68 | ES-33 day35 m2_004 | 1000x1100 | 11685 | 34140 | 97963 |
| 35 dpi | 23 | 69 | ES-33 day35 m2_005 | 1000x1100 | 10521 | | |
| 35 dpi | 24 | 70 | ES-33 day35 m3_001 | 1000x1100 | 6778 | | |
| 35 dpi | 24 | 71 | ES-33 day35 m3_002 | 1000x1100 | 10879 | 29118 | |
| 35 dpi | 24 | 72 | ES-33 day35 m3_003 | 1100x1000 | 11461 | | |
| Control | 25 | 73 | ES-33 day40 CONTROL m1_001 | 1000x1100 | 4958 | | |
| Control | 25 | 74 | ES-33 day40 CONTROL m1_002 | 1100x1000 | 11953 | 29643 | |
| Control | 25 | 75 | ES-33 day40 CONTROL m1_003 | 1000x1100 | 12732 | | |
| Control | 26 | 76 | ES-33 day40 CONTROL m2_001 | 1000x1100 | 10844 | | |
| Control | 26 | 77 | ES-33 day40 CONTROL m2_002 | 1100x1000 | 7300 | 27851 | 72316 |
| Control | 26 | 78 | ES-33 day40 CONTROL m2_003 | 1100x1000 | 9707 | | |
| Control | 27 | 79 | ES-33 day40 CONTROL m3_001 | 1000x1100 | 5411 | | |
| Control | 27 | 80 | ES-33 day40 CONTROL m3_002 | 1000x1100 | 7131 | 14822 | |
| Control | 27 | 81 | ES-33 day40 CONTROL m3_003 | 1000x1100 | 2280 | | |
| 40 dpi | 28 | 82 | ES-33 day40 m1_001 | 1000x1100 | 10605 | | |
| 40 dpi | 28 | 83 | ES-33 day40 m1_002 | 1100x1000 | 11694 | 30826 | |
| 40 dpi | 28 | 84 | ES-33 day40 m1_003 | 1000x1100 | 8527 | | |
| 40 dpi | 29 | 85 | ES-33 day40 m2_001 | 1000x1100 | 5432 | | |
| 40 dpi | 29 | 86 | ES-33 day40 m2_002 | 1100x1000 | 5920 | 15007 | 79989 |
| 40 dpi | 29 | 87 | ES-33 day40 m2_003 | 1100x1000 | 3655 | | |
| 40 dpi | 30 | 88 | ES-33 day40 m3_002 | 1100x1000 | 10197 | | |
| 40 dpi | 30 | 89 | ES-33 day40 m3_003 | 1100x1000 | 11532 | 34156 | |
| 40 dpi | 30 | 90 | ES-33 day40 m3_006 | 1100x1000 | 12427 | | |

**Table 3. Markers used for Identification of cell types via IMC – mouse.**

| **Broad cell types** | **Specific cell type** | **Markers** |
|---|---|---|
| Epithelial cells | | ICSK1 |
| | Proliferating Crypt cells | Ki-67, CD117 |
| | Crypt cells | Ki-67$_{low}$ |
| | Epithelial | - |
| | Shedded cells | - |
| Stroma | | - |
| | Smooth muscle cells and nerve fibers | Vimentin, Desmin, Collagen, VEGFR1, alphaSMA, GFAP |
| Endothelial | | CD31 |
| | VEC | VEGFR2, PV-1, VE-Cadherin |
| | Madcam-1+ VEC | PV-1, VE-Cadherin, Madcam-1 |
| | LEC | Lyve-1 |
| Immune cells | | CD45 |
| | Macrophage subset 1 | CD11b, CD206, CD163 |
| | Macrophage subset 2 and IELs | CD11b, F4/80, CD103, CD68, CD8 |
| | Macrophage subset 3 | CD11b, Lyve-1, CD206 |
| | CD8 | CD8, CD3, CD103 |
| | Immune cell patch | CD11c, Ki-67, CD54, CD5, CD4 |
| Undefined | | - |

**Table 4. B cell related genes removed from further analysis.**

| Ensemble gene id | External gene name | Description |
|---|---|---|
| ENSMUSG00000068105 | Tnfrsf13c | tumor necrosis factor receptor superfamily, member 13c |
| ENSMUSG00000061311 | Rag1 | recombination activating 1 |
| ENSMUSG00000049717 | Lig4 | ligase IV, DNA, ATP-dependent |
| ENSMUSG00000042474 | Fcmr | Fc fragment of IgM receptor |
| ENSMUSG00000040592 | Cd79b | CD79B antigen |
| ENSMUSG00000037922 | Bank1 | B cell scaffold protein with ankyrin repeats 1 |
| ENSMUSG00000032864 | Rag2 | recombination activating gene 2 |
| ENSMUSG00000032053 | Pou2af1 | POU domain, class 2, associating factor 1 |
| ENSMUSG00000030724 | Cd19 | CD19 antigen |
| ENSMUSG00000030577 | Cd22 | CD22 antigen |
| ENSMUSG00000030468 | Siglecg | sialic acid binding Ig-like lectin G |
| ENSMUSG00000029082 | Bst1 | bone marrow stromal cell antigen 1 |
| ENSMUSG00000027985 | Lef1 | lymphoid enhancer binding factor 1 |
| ENSMUSG00000027347 | Rasgrp1 | RAS guanyl releasing protein 1 |
| ENSMUSG00000024673 | Ms4a1 | membrane-spanning 4-domains, subfamily A, member 1 |
| ENSMUSG00000024353 | Mzb1 | marginal zone B and B1 cell-specific protein 1 |
| ENSMUSG00000020474 | Polm | polymerase (DNA directed), mu |
| ENSMUSG00000018168 | Ikzf3 | IKAROS family zinc finger 3 |
| ENSMUSG00000017652 | Cd40 | CD40 antigen |
| ENSMUSG00000014453 | Blk | B lymphoid kinase |
| ENSMUSG00000008193 | Spib | Spi-B transcription factor (Spi-1/PU.1 related) |
| ENSMUSG00000003379 | Cd79a | CD79A antigen (immunoglobulin-associated alpha) |

**Table 5. Results of PERMANOVA analysis from 16S rRNA analysis.**

| Sample type | Comparison | unweighted Unifrac | |
|---|---|---|---|
| | | **R2** | **Pseudo-F** |
| Complete | Control_vs_Day15 | **0.138** | **0.024** |
| | Control_vs_Day26 | **0.2152** | **0.001** |
| | Control_vs_Day35 | **0.1257** | **0.086** |
| | Control_vs_Day50 | **0.1381** | **0.028** |
| | Day15_vs_Day26 | **0.2943** | **0.029** |
| | Day15_vs_Day35 | **0.2662** | **0.029** |
| | Day15_vs_Day50 | **0.3277** | **0.02** |
| | Day26_vs_Day35 | **0.2455** | **0.02** |
| | Day26_vs_Day50 | **0.3087** | **0.034** |
| | Day35_vs_Day50 | **0.1995** | **0.075** |
| IgG | Control_vs_Day15 | **0.128** | **0.038** |
| | Control_vs_Day26 | **0.1457** | **0.053** |
| | Control_vs_Day35 | **0.1357** | **0.03** |
| | Control_vs_Day50 | **0.1376** | **0.04** |
| | Day15_vs_Day26 | **0.2947** | **0.057** |
| | Day15_vs_Day35 | **0.2373** | **0.03** |
| | Day15_vs_Day50 | **0.241** | **0.036** |
| | Day26_vs_Day35 | **0.285** | **0.106** |
| | Day26_vs_Day50 | **0.2316** | **0.146** |
| | Day35_vs_Day50 | **0.1915** | **0.027** |
| Control | IgG_vs_Complete | **0.2219** | **0.001** |
| Day15 | IgG_vs_Complete | **0.3729** | **0.019** |
| Day26 | IgG_vs_Complete | **0.112** | **0.834** |
| Day35 | IgG_vs_Complete | **0.3044** | **0.025** |
| Day50 | IgG_vs_Complete | **0.2807** | **0.025** |

**Table 6. Imaging Mass Cytometry panel – human.**

| Mass | Metal | Target | Clone | DF |
|---|---|---|---|---|
| 89 | Y | aSMA | 1A4 | 200 |
| 141 | Pr | CD38 | EPR4106 | 200 |
| 142 | Nd | CD19 | 6OMP31 | 200 |
| 144 | Nd | PIGR | 007 | 100 |
| 145 | Nd | t-bet | D6N8B | 50 |
| 146 | Nd | CD8a | C8/144B | 800 |
| 147 | Sm | CD163 | EDHu-1 | 100 |
| 148 | Nd | CD14 | EPR3653 | 100 |
| 149 | Sm | CD11b | EPR1344 | 300 |
| 150 | Nd | Ki-67 | B56 | 400 |
| 151 | Eu | CD31 | EPR3094 | 100 |
| 152 | Sm | CD45 | D9M8I | 100 |
| 154 | Sm | CD11c | 3.9 | 50 |
| 155 | Gd | FoxP3 | PCH101 | 200 |
| 156 | Gd | CD4 | EPR6855 | 100 |
| 158 | Gd | E-Cadherin | 2.4E10 | 200 |
| 159 | Tb | CD68 | KP1 | 1000 |
| 160 | Gd | IL17A | AF-317-NA | 200 |
| 161 | Dy | CD20 | 2H7 | 50 |
| 162 | Dy | CD144 (VE-cadherin) | 16B1 | 100 |
| 163 | Dy | gata3 | EPR16651 | 50 |
| 164 | Dy | CD62L | PA5-35327 | 100 |
| 167 | Er | CD103 | BLR171J | 100 |
| 168 | Er | CD127 | EPR2955(2) | 200 |
| 169 | Tm | Collagen Type I | Polyclonal | 500 |
| 170 | Er | CD3 | Polyclonal | 200 |
| 171 | Yb | CD27 | EPR8569 | 200 |
| 172 | Yb | cKit /CD117 | YR145 | 200 |
| 173 | Yb | CD45RO | UCHL1 | 200 |
| 174 | Yb | Podoplanin | NC08 | 200 |
| 175 | Lu | CD56 | 07-5603 | 100 |
| 176 | Yb | Histone 3 | D1H2 | 500 |
| 191/193 | Ir | DNA | NA | 400 |

**Table 7. Samples list with cell numbers of via IMC analysed ileal tissue – human.**

| Type | Condition | Patient ID | Sample ID | ROI | ROI size | Cells per ROI | Total cells per Patient/Condition | Total cells per group |
|---|---|---|---|---|---|---|---|---|
| Epithelial | early_RA | 14 | 59 | ROI009_Patient 14_2 | 1000 x 1000 | 3539 | 0 | 73076 |
| | early_RA | 16 | 53 | ROI008_Patient 16_1 | 1000 x 1000 | 6435 | 6435 | |
| | early_RA | 20 | 2 | ROI001_Patient 20_2 | 1000 x 1000 | 4547 | 4547 | |
| | early_RA | 22 | 20 | ROI003_Patient 22_1 | 1000 x 1000 | 6361 | 6361 | |
| | early_RA | 24 | 5 | ROI001_Patient 24 | 1000 x 1000 | 3822 | 10323 | |
| | early_RA | 24 | 26 | ROI003_patient 24_1 | 1000 x 1000 | 6501 | | |
| | early_RA | 29 | 14 | ROI002_Patient 29_2 | 1000 x 1000 | 3583 | 3583 | |
| | early_RA | 42 | 24 | ROI003_Patient 42_1 | 1692 x 709 | 5919 | 11503 | |
| | early_RA | 42 | 56 | ROI008_Patient 42_2 | 1000 x 1000 | 5584 | | |
| | early_RA | 43 | 32 | ROI004_Patient 43_1 | 1000 x 1000 | 5919 | 12328 | |
| | early_RA | 43 | 39 | ROI005_Patient 43_2 | 1000 x 927 | 6409 | | |
| | early_RA | 46 | 33 | ROI004_Patient 46_2 | 1000 x 1000 | 4432 | 4432 | |
| | early_RA | 49 | 57 | ROI008_Patient 49_2 | 1000 x 1000 | 6869 | 6869 | |
| | healthy | 3 | 23 | ROI003_Patient 3_1 | 1000 x 1000 | 7731 | 14006 | 106729 |
| | healthy | 3 | 55 | ROI008_Patient 3_2 | 1000 x 1000 | 6275 | | |
| | healthy | 6 | 8 | ROI001_Patient 6_1 | 1000 x 1000 | 5290 | 11299 | |
| | healthy | 6 | 68 | ROI010_Patient 6_2 | 1000 x 1000 | 6009 | | |
| | healthy | 17 | 35 | ROI005_Patient 17_2 | 900 x 1100 | 6311 | 11168 | |
| | healthy | 17 | 42 | ROI006_Patient 17_1 | 1000 x 1000 | 4857 | | |
| | healthy | 19 | 19 | ROI003_Patient 19_2 | 1000 x 1000 | 5635 | 11290 | |
| | healthy | 19 | 28 | ROI004_Patient 19_1 | 900 x 1100 | 5655 | | |
| | healthy | 23 | 12 | ROI002_Patient 23_2 | 1000 x 1000 | 5217 | 5217 | |
| | healthy | 25 | 13 | ROI002_Patient 25_1 | 1000 x 1000 | 7874 | 16235 | |
| | healthy | 25 | 21 | ROI003_Patient 25_2 | 1000 x 1000 | 8361 | | |
| | healthy | 27 | 30 | ROI004_Patient 27_1 | 1000 x 1000 | 7481 | 7481 | |
| | healthy | 39 | 75 | ROI012_Patient 39_2 | 1000 x 1000 | 5001 | 5001 | |

| | | | | | | | | |
|---|---|---|---|---|---|---|---|---|
| | healthy | 44 | 45 | ROI006_Patient 44_1 | 1088 x 876 | 397 0 | 10543 | |
| | healthy | 44 | 50 | ROI007_Patient 44_2 | 1054 x 926 | 657 3 | | |
| | healthy | 48 | 40 | ROI005_Patient 48_1 | 1000 x 1000 | 843 9 | 14489 | |
| | healthy | 48 | 46 | ROI006_Patient 48_2 | 1000 x 1000 | 605 0 | | |
| | IBD | 13 | 73 | ROI012_Patient 13_1 | 1000 x 1000 | 584 6 | 5846 | 3959 1 |
| | IBD | 21 | 11 | ROI002_Patient 21_2 | 1000 x 1000 | 560 7 | 5607 | |
| | IBD | 33 | 49 | ROI007_Patient 33_1 | 1000 x 1000 | 731 5 | 7315 | |
| | IBD | 40 | 61 | ROI009_Patient 40_1 | 1000 x 1000 | 526 4 | 5264 | |
| | IBD | 45 | 7 | ROI001_Patient 45_1 | 1000 x 1000 | 872 6 | 15559 | |
| | IBD | 45 | 16 | ROI002_Patient 45_2 | 800 x 1200 | 683 3 | | |
| | RA | 1 | 43 | ROI006_Patient 1_1 | 1000 x 1000 | 692 3 | 14296 | 9440 1 |
| | RA | 1 | 48 | ROI007_Patient 1_2 | 1000 x 1000 | 737 3 | | |
| | RA | 2 | 74 | ROI012_Patient 2_2 | 1000 x 1000 | 578 1 | 5781 | |
| | RA | 5 | 17 | ROI002_Patient 5_1 | 1000 x 1000 | 622 1 | 6221 | |
| | RA | 7 | 72 | ROI011_Patient 7_1 | 1000 x 1000 | 711 5 | 15598 | |
| | RA | 7 | 76 | ROI012_Patient 7_2 | 1000 x 1000 | 848 3 | | |
| | RA | 8 | 64 | ROI009_Patient 8_1 | 1000 x 1000 | 700 7 | 12706 | |
| | RA | 8 | 69 | ROI010_Patient 8_2 | 1000 x 1000 | 569 9 | | |
| | RA | 9 | 58 | ROI008_Patient 9_2 | 1000 x 1000 | 694 1 | 6941 | |
| | RA | 10 | 34 | ROI005_Patient 10_1 | 1000 x 1000 | 590 4 | 11176 | |
| | RA | 10 | 41 | ROI006_Patient 10_2 | 1000 x 1000 | 527 2 | | |
| | RA | 11 | 27 | ROI004_Patient 11_2 | 900 x 1100 | 289 3 | 2893 | |
| | RA | 32 | 38 | ROI005_Patient 32_1 | 1000 x 1000 | 653 2 | 12654 | |
| | RA | 32 | 44 | ROI006_Patient 32_2 | 1000 x 1000 | 612 2 | | |
| | RA | 37 | 66 | ROI010_Patient 37_2 | 1000 x 1000 | 613 5 | | |
| Immune | early_RA | 14 | 65 | ROI010_Patient 14_1 | 1000 x 1000 | 678 7 | 6787 | 7423 2 |
| | early_RA | 20 | 10 | ROI002_Patient 20_1 | 1000 x 1000 | 926 9 | 9269 | |
| | early_RA | 22 | 29 | ROI004_Patient 22_2 | 1000 x 1000 | 128 14 | 12814 | |
| | early_RA | 29 | 6 | ROI001_Patient 29_1 | 1000 x 1000 | 711 4 | 7114 | |

| | | | | | | | | |
|---|---|---|---|---|---|---|---|---|
| | early_RA | 31 | 31 | ROI004_Patient31_2 | 1000 x 1000 | 10777 | 10777 | |
| | early_RA | 46 | 25 | ROI003_Patient46_1 | 1000 x 1000 | 11297 | 11297 | |
| | early_RA | 49 | 51 | ROI007_Patient49_1 | 1000 x 1000 | 16174 | 16174 | |
| | healthy | 27 | 36 | ROI005_Patient27_2 | 1000 x 1000 | 15830 | 15830 | 15830 |
| | IBD | 12 | 1 | ROI001_Patient12_1 | 1000 x 1000 | 9948 | 23973 | 47052 |
| | IBD | 12 | 9 | ROI002_Patient12_2 | 1000 x 1000 | 14025 | | |
| | IBD | 21 | 3 | ROI001_Patient21_1 | 1000 x 1000 | 11311 | 11311 | |
| | IBD | 33 | 54 | ROI008_Patient33_2 | 1000 x 1000 | 11768 | 11768 | |
| | RA | 2 | 37 | ROI005_Patient2_1 | 1200 x 1100 | 12268 | | 12268 |
| Immune/Epithelial | early_RA | 16 | 47 | ROI007_Patient16_2 | 900 x 1100 | 6088 | 6088 | 17568 |
| | early_RA | 31 | 22 | ROI003_Patient31_1 | 1000 x 1000 | 11480 | 11480 | |
| | healthy | 23 | 4 | ROI001_Patient23_1 | 1000 x 1000 | 6962 | 6962 | 15337 |
| | healthy | 39 | 71 | ROI011_Patient39_1 | 1000 x 1000 | 8375 | 8375 | |
| | IBD | 13 | 70 | ROI011_Patient13_2 | 1000 x 1000 | 9071 | 9071 | 9071 |
| | RA | 5 | 63 | ROI009_Patient5_2 | 1000 x 1000 | 7869 | 7869 | 21120 |
| | RA | 9 | 52 | ROI007_Patient9_1 | 1000 x 1000 | 8236 | 8236 | |
| | RA | 11 | 18 | ROI003_Patient11_1 | 900 x 1100 | 5015 | 5015 | |

**Table 8. Markers used for Identification of cell types via IMC – human.**

| Broad cell types | Specific cell type | Markers |
|---|---|---|
| Epithelial cells | - | E-cadherin |
| | Epithelial | - |
| | Crypt cells | Ki-67, PIGR |
| | Goblet | - |
| Stroma | - | Collagen I |
| | Fibroblast | - |
| Endothelial | - | CD31 |
| | LEC | Lyve-1 |
| | VEC | CD144 |
| Immune cells | - | CD45 |
| | CD8a CD103 | CD3, CD8a, CD103 |
| | CD8a | CD3, CD8a |
| | CD4 | CD3, CD4 |
| | Macrophage | CD163 |
| | Plasma cell | CD38, CD27 |
| | $cKit^{+}$ | $cKit^{+}$ |
| | Bcell | CD19 |
| | Proliferating immune | Ki-67 |
| Undefined | - | - |

**Table 9. Abbreviation table.**

| Abbreviation | Definition |
|---|---|
| anti-citrullinated protein antibodies | ACPA |
| anti-modified protein antibodies | AMPA |
| Benjamini-Hochberg | BH |
| cellular neighbourhood | CN |
| chicken type II collagen | CII |
| collagen-induced arthritis | CIA |
| Complete Freund's Adjuvant | CFA |
| days post-immunization | dpi |
| Deutsche Forschungsgemeinschaft | DFG |
| differentialy expressed genes | DEG |
| false discovery rate | FDR |
| gene ontology | GO |
| imaging mass cytometry | IMC |
| immunoglobulin G | IgG |
| inflammatory bowel disease | IBD |
| interquartile range | IQR |
| intraepithelial lymphocytes | IEL |
| lipopolysaccharide-binding protein | LBP |
| principal component analysis | PCA |
| principal coordinate | PCo |
| principal coordinate analysis | PCoA |
| region of interests | ROI |
| rheumatoid arthritis | RA |
| short-chain fatty acids | SCFA |
| smooth muscle cell | SMC |
| soluble CD14 | sCD14 |
| systemic lupus erythematosus | SLE |
| type 1 interferon | IFN |
| vascular endothelial cells | VEC |
