## Supplementary Methods for "Microbiota-specific serum IgG links gut and joints through immune–endothelial crosstalk in arthritis"

### *In vivo* imaging of colonic endothelial leakiness

In vivo assessment of colonic vascular permeability was performed as described previously (Langer et al., 2019) in collaboration with AG Stürzl at the Translational Research Center (TRC, Erlangen, Germany). To evaluate endothelial leakage, fluorescein isothiocyanate (FITC)-dextran (70 kDa; Sigma-Aldrich) was administered as a vascular tracer. For vessel contrast, Bandeiraea simplicifolia lectin was conjugated to Cy5 using a monoreactive N-hydroxysuccinimide ester (Roche). All fluorescent reagents were stored at 4 °C in the dark and freshly prepared on the day of imaging. FITC-dextran was dissolved in sterile phosphate-buffered saline (PBS) at 10 mg/ml. For combined vessel visualization and permeability detection, FITC-dextran (20 mg/ml) was mixed 1:1 with Cy5-labeled lectin to yield a final injection volume of 150 µL per mouse. Lectin was diluted to a final dose of 100 µg per mouse (for example, 18.72 µL of a 5.34 mg/ml stock combined with 75 µL PBS, then mixed with 75 µL FITC-dextran solution).

Mice were anesthetized via intraperitoneal injection of a ketamine/xylazine mixture (120 mg/kg ketamine and 8 mg/kg xylazine; stocks: 100 mg/ml and 20 mg/ml, respectively). For a 25 g mouse, this corresponded to 30 µL ketamine, 10 µL xylazine, and 160 µL PBS (total = 200 µL). Anaesthesia depth was confirmed by loss of pedal reflex and adjusted as needed. The tracer mixture (150 µL) was administered intravenously via retrobulbar injection.

After tracer circulation, a midline laparotomy was performed to expose the colon. A short colonic segment was excised, opened longitudinally, and gently rinsed with sterile PBS. Residual blood was carefully removed without disrupting the mucosal surface. The prepared tissue was mounted luminal side up on a custom imaging platform and analysed by laser scanning confocal microscopy (Leica TCS SPE, LAS LAF software). FITC-dextran extravasation served as an indicator of vascular leakage.

After imaging, mice were euthanized by cervical dislocation. Vascular permeability was quantified by determining the proportion of FITC-positive crypts within a defined region of interest (ROI) containing fully imaged crypts.

### Imaging mass cytometry (IMC)

Preparation of murine ileal tissue: Following euthanasia by cervical dislocation, the distal third of the small intestine (ileum) was excised. Luminal contents were gently removed, and the tissue was rinsed with ice-cold PBS. Residual PBS was carefully blotted before embedding in OCT compound. To prepare intestinal rolls, the ileum was rolled longitudinally, placed into a cryomold partially filled with OCT (approximately one-third of its volume), and positioned on dry ice to initiate solidification. The intestinal roll was arranged within the mold, additional OCT was added until fully covered, and the block was solidified on dry ice before storage at –80 °C. Cryosections of 5 µm thickness were prepared using a cryostat and mounted on Epredia™ SuperFrost Plus™ adhesion microscope slides (Thermo Fisher Scientific, Waltham, USA). Sections were air-dried for 30 min at room temperature and stored again at –80 °C until further processing. On the day of staining, tissue sections were fixed with freshly prepared 4 % paraformaldehyde (PFA) for 5 min at room temperature, rehydrated for 5 min in PBS, and washed for 5 min in wash buffer (PBS supplemented with 0.05% Tween and 1% bovine serum albumin (BSA)). A hydrophobic barrier was drawn around each section to confine reagents. To block nonspecific antibody binding, 100 µL of SuperBlock™ (Thermo Fisher Scientific, Waltham, USA) solution was applied for 1 h at room temperature. Excess blocking solution was gently tapped off, and 50 µL of antibody mixture (**Supplementary Table 1**), diluted in Dako REAL Antibody Diluent (Agilent, Santa Clara, USA), was added and incubated overnight at 4 °C in a humidified chamber. Sections were then washed three times for 5 min in wash buffer. Subsequently, slides were incubated with 100 µL Cell-ID™ Intercalator-Ir (Standard BioTools, San Francisco, USA; 1:400 in PBS) for 30 min at room temperature, rinsed once, and washed twice for 5 min with wash buffer. Slides were finally rinsed in Milli-Q water and air-dried in a fume hood. For each mouse, three regions of interest (ROIs) of 1000 µm × 1100 µm were acquired on the Hyperion Imaging System (Standard BioTools, San Francisco, USA) using a laser power of 2 and an ablation frequency of 200 Hz.

Data analysis of murine IMC data: 16bit single stack .tiff files were exported with MCD viewer and the IMC Denoise pipeline (1) was used to improving the quality of the IMC data. The pipeline includes differential intensity map-based restoration (DIMR) to remove hot pixels and deep self-supervised noise filtering (DeepSNF) to reduce noise for all used channels. Stack images in .tiff format were created using ImageJ. The Steinbock open-source software was used for cell segmentation with the Deepcell deep learning library (nuclei: DNA; cytoplasm: ICSK1 and ICSK2), quantitative feature extraction and data export (2). Files were imported in to R (v 4.3.3) and analyzed using the IMCDataAnalysis pipeline by Windhager *et. al* (2023) (2). In short, the read_steinbock function from the IMCtools package (2) was used to load the single cell data in R, and a spatial experiment (spe) object was generated. Counts were transformed calculating the inverse hyperbolic sine using the asinh function, dividing the values through 0.2 and saving the values as expression values. Use channels (excluded markers: DNA, Histone, Ly6G, CD127, CD69, ICSK2, CD27, ICSK3, RORgammaT, FoxP3, CD19, T-bet) were defined for dimensionality reduction, subsequent batch effect correction and clustering. Cells with an area below 4 were excluded from further analysis. Next, with the runUMAP function, a linear dimensionality reduction method to project cells from a high-dimensional down to a low-dimensional space, was performed with the expression values using the scater package (3, 4). Harmony correction (ncomponents = 30) of the expression values was performed to correct for batch effects (5). The Rphenograph (k = 50) function was used to cluster harmony corrected data (6, 7). Cell types were defined by spatial location and marker expression. For cell – cell interaction analysis, using the buildSpatialGraph function, interaction was calculated using a Delaunay triangulation based graph construction with max_dist set to 20 (8). Using the aggregate Neighbors function, six different cellular neighborhoods (CNs) were defined based on the fraction of a certain cell type among its 20 nearest neighbours. The Number of Interactions between cell types in one CN per ROI were normalized on the cell number in the CN per ROI ($n_{CNinteractions}/n_{CNcellcount}$ *100). Normalized interactions with mean (all groups) normalized interactions below 90 were excluded.

Preparation of human tissue for IMC: Biopsies were fixated in a 4% PFA solution and dehydrated in a routine process overnight and embedded in paraffin. Sections for IHC-stainings were taken at 3.5-4 µm and mounted on Super Frost Plus microscope slides. Formalin-fixed, paraffin-embedded (FFPE) tissue sections were stained for IMC according to an optimized protocol adapted from Standard Biotools (IMC Staining Protocol for FFPE Sections, 2022). Tissue sections mounted on glass slides were baked at 60 °C for 2 hours to remove residual paraffin. Slides were then dewaxed in fresh xylene for 20 min followed by rehydration through a graded ethanol series (100 %, 95 %, 80 %, and 70 %, 5 min each). Antigen retrieval was performed by incubating slides in preheated antigen retrieval solution (pH 9.1×, prepared from a 10× stock; Agilent, Santa Clara, USA) at 96 °C for 30min. Following incubation, slides were cooled to 70 °C for 10 min. Slides were then washed in MiliQ Water for 5 min followed by washing step in PBS for 10 min, both with gentle agitation. Tissue sections were encircled with a hydrophobic barrier using a barrier pen and blocked with 3 % BSA in PBS for 45 min at room temperature in a humidified chamber. The metal-conjugated antibody mix (**Supplementary Table 6**), diluted in Dako REAL Antibody Diluent (Agilent, Santa Clara, USA), was applied to the tissue sections, which were then incubated overnight at 4 °C in a humidified chamber. Following primary antibody incubation, slides were washed twice in 0.2 % Triton™ X-100 Surfact-Amps™ (Thermo Fisher Scientific, Waltham, USA) in PBS for 8min each and then twice in PBS for 8 min each, all with gentle agitation. Nuclear staining was performed using Cell-ID™ Intercalator-Ir (Standard BioTools, San Francisco, USA; 1:400 in PBS) for 30 min at room temperature. Slides were washed in MiliQ Water for 5 min with gentle agitation and then air-dried for at least 20 min at room temperature prior to IMC data acquisition on the Hyperion Imaging System Hyperion System with a laser power of 2 and an ablation frequency of 200 Hz.

Data analysis of human IMC: 16bit single stack .tiff files were exported with MCD viewer and the IMC Denoise pipeline (1) was used to improving the quality of the IMC data. The pipeline includes DIMR to remove hot pixels and DeepSNF to reduce noise for all used channels. Stack images in .tiff format were created using ImageJ. The Steinbock open-source software was used for cell segmentation with the Deepcell deep learning library (nuclei: DNA; cytoplasm: ICSK1 and ICSK2), quantitative feature extraction and data export (2). Files were imported in to R (v 4.3.3) and analyzed using the IMCDataAnalysis pipeline by Windhager *et. al* (2023) (2). In short, the read_steinbock function from the IMCtools package (2) was used to load the single cell data in R, and a spe object was generated. Counts were transformed calculating the inverse hyperbolic sine using the asinh function, dividing the values through 0.2 and saving the values as expression values. Use channels (excluded markers: DNA, Histone, CD11b, Gata3, CD11c, CD20, Tbet, CD127) were defined for dimensionality reduction, subsequent batch effect correction and clustering. Cells with an area below 5 were excluded from further analysis. Next, using the runUMAP function, a linear dimensionality reduction projecting cells from a high-dimensional down to a low-dimensional space, was performed with the expression values using the scater package (3, 4). Harmony correction (ncomponents = 30) of the expression values was performed to correct for batch effects (5). The Rphenograph (k = 50) function was used to cluster harmony corrected data (6, 7). Cell types were defined by spatial location and marker expression. For cell – cell interaction analysis, using the buildSpatialGraph function, interaction was calculated using a Delaunay triangulation based graph construction with max_dist set to 20 (8). Using the aggregate Neighbors function, six different CNs were defined based on the fraction of a certain cell type among its 20 nearest neighbors. The Number of Interactions between cell types in one CN per ROI were normalized on the cell number in the CN per ROI ($n_{CNinteractions}/n_{CNcellcount}*1000$).

**Bulk RNA sequencing analysis of murine endothelial cells**

Isolation of intestinal endothelial cells: Intestinal endothelial cells were isolated as previously described (9) with minor modifications. Mice were euthanized using $CO_2$ asphyxiation and perfused with 20 ml of ice-cold PBS to flush erythrocytes from the vasculature. The entire small intestine was excised, and the proximal section was cut approximately 4 cm distal to the stomach, while the distal section was cut proximal to the caecum to collect the jejunum and ileum. The intestinal lumen was flushed with 20 ml ice-cold PBS, opened longitudinally, and gently cleared of mucus using a cover glass, avoiding disruption of villi. The tissue was cut into ~0.5 cm fragments and incubated in pre-digestion solution (PBS containing 10 mM EDTA and 0.4% v/v endothelial cell growth supplement, Promocell, PromoCell, Heidelberg, Germany) with gentle agitation at 37 °C for 20 min to detach epithelial cells. The suspension was briefly vortexed (30 s, maximum speed) and the supernatant discarded. Samples were washed three times in 20 ml ice-cold PBS with vortexing until the supernatant was clear.

The remaining tissue was finely minced and transferred into gentleMACS™ C Tubes (Miltenyi Biotec B.V. & Co. KG, Bergisch Gladbach, Germany) containing 10 ml digestion buffer (0.2% FCS, Gibco, Dublin, Ireland; 3 mg/ml Collagenase IV, Sigma, St. Louis, USA; 0.2 mg/ml DNAse I, Sigma, St. Louis, USA; 2mM $CaCl_2$, Roth, Karlsruhe, Germany; 0.4% Endothelial Cell Growth Supplement/ Heparin, PromoCell, Heidelberg, Germany) in Dulbecco's Modified Eagle Medium (DMEM, Thermo Fisher Scientific, Waltham, USA). Tissue was dissociated using the pre-programmed "brain" protocol on the MACS Tissue Dissociator (Miltenyi Biotec B.V. & Co. KG, Bergisch Gladbach, Germany), followed by enzymatic digestion on a thermo-shaker at 37 °C, 200 rpm, for 20 min. The dissociation step was repeated twice, and digestion was stopped by adding 10 ml DMEM supplemented with 10% FCS. Cell suspensions were centrifuged (300 × g, 5 min, 4 °C), resuspended in 20 ml PBS containing 1% 100x Penicillin/Streptomycin (Thermo Fisher Scientific, Waltham, USA), and filtered sequentially through 100 µm, 70 µm, and 40 µm strainers.

For immunostaining, cells were pelleted (300 × g, 5 min, 4 °C), resuspended in antibody mix containing anti-CD45 (1:400, Biolegend, 304054) and anti-CD31 (1:200, Thermo Fisher scientific, 17-0311-80) solved in FACS buffer, and stained for 15 min on ice. Staining was terminated with FACS buffer (PBS containing 2% FCS, Gibco, Dublin, Ireland; 2 mM EDTA (0.5M), Merck, Darmstadt, Germany) containing DAPI (0.625 µg/ml). After washing and centrifugation, cells were resuspended in 350 µL FACS buffer, filtered through a 40 µm strainer, and prepared for flow cytometric sorting. Endothelial cells were identified as $DAPI^-$ $CD45^-$, $CD31^+$ events. Endothelial cells were sorted with a FACS Aria III with an 85 µm nozzle. In total, a minimum of 65.000 intestinal endothelial cells were sorted and collected in 350 µl RLT buffer (Qiagen, Venlo, Netherlands) and stored at -80 °C until RNA extraction.

Isolation of bone endothelial cells: Mice were sacrificed by cervical dislocation and legs were removed to obtain femur and tibiae from both sides. After removing the knee, bones were crushed in 2 ml ice-cold PBS containing 4 % FCS, using a mortar and a pestle. The obtained supernatant was filtered through a 100 µm pre-wetted cell strainer into a 50 ml falcon. This was repeated two more times and the filter was washed with 2 ml ice-cold PBS containing 4 % FCS to reach a total volume of 10 ml. After centrifugation (400 rcf, 5min, 4 °C) the supernatant was discarded. The pellet was solved in 1 ml PBS and 9 ml deionized water was added for erythrocyte lysis for 30 sec. To stop the reaction, 1 ml 10x PBS was added and centrifuged (400 rcf, 5min, 4 °C). The pellet was resuspended in 1 ml biotinylated antibody mix containing anti-CD45 (1:500, Thermo Fisher scientific, 13-0451-82) and anti-Ter-119 (1:200, Biolegend, 116204) in FACS buffer and incubated for 30 min at 4°C on a shaker. Cells were washed with 1 ml of FACS buffer and centrifuged (400 rcf, 5min, 4 °C). The supernatant was removed, the pellet was resuspended in 1 ml of PBS containing 4 % FCS and 70 µl Dynabeads™ Biotin Binder (Thermo Fisher scientific, Waltham, USA) and incubated in a rotary mixer at 4 °C for 20 min. Thereafter, cells were Incubate in magnet for 5 min. Non-bound cells were transferred in a new FACS tube and 2 ml of FACS buffer were added to the cells in the magnet. After another 5 min incubation in the magnet, non-bound cells were added to the previous cells. Cells were centrifuged (400 rcf, 5 min, 4 °C), the supernatant discarded, the pellet resuspended in 1 ml staining antibody mix containing Lineage antibody cocktail (containing anti CD3, Ly-6G, CD11b, CD45R, Ter-119; 1:1000, Biolegend, 133311), anti-Sca-1 (1:200, Thermo Fisher scientific, 12-5851-82), anti-CD31 (1:600, Thermo Fisher scientific, 17-0311-80) and anti-Endomucin (1:200, Thermo Fisher scientific, 12-5851-82) solved in FACS buffer and incubated for 30 min at 4°C. To wash the cells, 1 ml of FACS buffer was added and the suspension was centrifuged (400 rcf, 5min, 4 °C). The supernatant was discarded and the pellet was stained with the LIVE/DEAD™ Fixable Aqua Dead Cell Stain Kit (1:800 In $H_2O$, Thermo Fisher scientific, Waltham, USA) and incubated for 15 min at RT. The staining was stopped by adding FACS buffer. After centrifugation (400 rcf, 5 min, 4 °C) cells were reconstituted in 350 µL FACS buffer, filtered through a 40 µm cell strainer and ready for the acquisition. Endothelial cells were identified as viable, Lineage$^-$, Sca-1$^-$, CD31$^+$ events. total, a minimum of 100,000 bone endothelial cells (viable Lin- Sca1- CD31+) were sorted in a 5 ml FACS tube containing 350 µl RLT buffer (Qiagen, Venlo, Netherlands) and stored at -80 °C until RNA extraction.

RNA extraction for mRNA sequencing analysis: For the bulk mRNA sequencing analysis of sorted intestinal and bone endothelial cells, total RNA was isolated using a RNeasy plus Micro Kit plus (Quiagen, Netherlands) according to the manufacturer's recommendations. Measurement of RNA concentration was performed with a NanoDrop 2000 (Thermo Fisher Scientific, Waltham, USA). First, 1 µl RNase free water was measured as a blank. Then, 1 µl of each sample was measured to determine RNA concentration and purity.

Bulk mRNA sequencing and data analysis: The Illumina RNA sequencing analysis was performed by Novogene. In brief, sequencing libraries were generated using NEBNext® Ultra TM RNA Library Prep Kit for Illumina® (NEB, USA) and sequenced on an Illumina platform. Raw data (raw reads) of FASTQ format were processed through fastp. Mapping of the processed data to the reference genome mus musculus (for intestinal: GRCm39/mm39; for bone: GRCm38/mm10) was performed using hisat2 (10). FeatureCounts was used for the quantification of the mapped reads (11). Raw mapped reads were processed in R (R version 4.3.3, Lucent Technologies) with DESeq2 (12), to determine differentially expressed genes and generate normalized read counts. Before processing the intestinal counts with DESeq2, raw counts were selected for protein coding genes and genes with zero transcripts were removed. For principal component analysis (PCA) normalized counts were log transformed. The top 200 genes with most variable genes contributing to principal component 1 (PC1) were used for pathway enrichment analysis using the clusterProfiler package (maxGSsize = 200, pAdjustMethod = Benjamini-Hochberg) (13). Heatmaps were generated using the pheatmap package (14). Differentially expressed genes (DEG) were defined as genes with a $padj < 0.05$ and log2 Fold change (FC) > |1.5| compared to controls and used for pathway enrichment analysis. Venn diagrams of DEGs were generated using the ggvenn package (15). Volcano plots were generated using the EnhancedVolcano package (16). Graphs were generated using the ggplot2 package (17). Deconvolution of the sorted bone cells was performed with the immunedeconv package using the deconvolute_mouse function (setting: mmcp_counter) of TPM converted counts and the reference genome GRCm38/mm10 (18, 19).

**Analysis of serum IgG bound fecal bacteria**

Preparation of stool for isolation of serum-IgG bound bacteria: For sorting of serum IgG bound faecal bacteria, faecal pellets were collected directly from mice (~100 mg, 3 pellets), placed in tubes with ceramic beads (Soft tissue homogenizing CK14 2 ml Lysing Kit, BioCat GmbH, Heidelberg, Germany) and incubated in 1 ml PBS per 100 mg faecal material for 1 h on ice. The samples were homogenized by vortexing until a homogenous solution has formed and centrifuged (50 rcf, 15 min, 4 °C) to remove large particles. 300 µl of the supernatant were transferred to another microcentrifuge tube and washed with 1 ml PBS containing 1% (w/v) BSA (staining buffer) and centrifuged for 5 min (8,000 rcf, 4°C) before resuspension in 1 ml staining buffer. After two additional washes, bacterial pellets were resuspended in 300 µl blocking buffer (staining buffer containing 20% normal rat serum) and incubated for 30 min on ice. Samples were washed with 1ml staining buffer and from each sample 50 µl were taken as a pooled unstained control. After one centrifugation step (800 rcf. 4 °C, 5 min), samples were stained with 100 µl staining buffer containing PE-conjugated Anti-Mouse IgA (1:12.5; Thermo Fisher scientific, 12-4204-82) and FITC-anti-igG (1:50, Biolegend, 405305) for 30 min on ice. Samples were washed with 1 ml staining buffer before 1:2 serum dilution was incubated with half of the volume of pre-stained stool samples overnight at 4°C at the shaker to identify to which bacteria serum IgG binds. After two washing steps, samples were stained with AF647-anti-igG (1:50, Biolegend, 405322) for 30 min on ice. As a negative control the other half of pre-stained stool was incubated with AF647-anti-igG (1:50, Biolegend, polyclonal) for 30 min on ice. After washing twice with 1 ml PBS (10,000 rcf, 5 min, 4 °C), the serum-IgG$^{high}$ (high AF647 signal) bacteria were sorted via FACS (Cytoflex SRT, Beckman Coulter, USA), and pre-sort samples were used as complete controls for 16S rRNA anaysis. In total, a minimum of 100,000 IgG$^{high}$ events were sorted in a 5 ml FACS tube, centrifuged (10,000 rcf, 5 min, 4 °C), the supernatant was removed and stored at -80 °C until DNA extraction.

<u>DNA extraction:</u> DNA was extracted using the PureLink™ Microbiome DNA Extraction kit (Thermo Fisher Scientific, Waltham, USA) according to the manufacturer's instructions.

16S rRNA Sequencing: The 16S rRNA Sequencing was performed by the core facility of the university clinic Erlangen. In short, 10 ng of stool genomic DNA was used for PCR amplification of the 16S ribosomal RNA V4 region using the prokaryotic primer pair 515F (5'-GTGYCAGCMGCCGCGGTAA-3') and 806R (5'-GGACTACNVGGGTWTCTAAT-3') containing barcodes on the forward primer 515F (https://earthmicrobiome.org/protocols-and-standards/16s/). PCR was performed using the NEBNext Q5 Hot Start HiFi PCR Master Mix (New England Biolabs, Frankfurt am Main, Germany) with 25 cycles. Amplicons were purified with AMPure XP beads (Beckmann Coulter GmbH, Krefeld, Germany), pooled in equimolar ratios, and sequenced by 2 × 151 paired-end sequencing on an Illumina MiSeq system (Illumina Inc., San Diego, USA). Raw FASTQ files were imported and processed in QIIME2 (v2022.2), where DADA2 was used for quality control, dereplication, and generation of amplicon sequence variants (ASVs). Taxonomic classification was performed against the SILVA small subunit database release 138 at 99% similarity cutoff. ASV and taxonomy tables were imported into R (v4.2) as phyloseq objects. Alpha diversity indices, including inverse Simpson diversity, were calculated after rarefaction to the smallest library size. Beta diversity was assessed by computing weighted and unweighted UniFrac distance matrices without log transformation. Differences in microbial community composition between groups were tested by permutational multivariate analysis of variance (PERMANOVA) using the adonis function in the vegan R package (v2.6.4). Differential abundance analysis of taxa was conducted using Linear Discriminant Analysis Effect Size (LEfSe) to identify biomarker taxa associated with experimental groups. Visualization of diversity metrics and ordination results was performed with the ggplot2 package.

**Quantification of commensal bacteria-reactive IgG using ELISA**

For measurement of commensal bacteria-reactive IgG, fecal bacteria were isolated from naive CIA mice and used to coat medium binding ELISA (Greiner AG, Kremsmünster, Austria). plates. Therefore, fecal bacteria from ~2-3 stool pellets were isolated by homogenization (vortex ~40 sec) in 1 ml sterile PBS and filtered through a 40 µm cell strainer. The filter was washed with 3 ml PBS and collected in the same tube. The bacteria were separated from debris/mouse cells by removing the pellet after centrifugation at 1,000 rcf for 5 min. The resulting supernatant was washed with sterile PBS twice by centrifuging for 1 min at 8,000 rcf. On the last wash, bacteria were re-suspended in 2 ml ice-cold PBS and sonicated on ice (3 x 30 sec, amplitude = 37 %). Samples were then centrifuged at 16,000 rcf for 10 min, and supernatants recovered for a crude commensal bacteria antigen preparation. For measurement of serum antibodies by ELISA, 5 µg/ml commensal bacteria antigen was coated on medium binding 96-well plates overnight (4 °C), washed extensively and blocked 1 h with 3 % BSA in PBS (300 µL/well). 100 µL sera were incubated overnight in doubling dilutions solved in 1 % BSA. Antigen-specific IgG etc. was detected incubating a goat anti-mouse IgG horseradish peroxidase (HRP) coupled antibody (1:8000, Southern Biotech, 1030-05) at RT. Plates were washed extensively before they were developed with EBioscience™ TMB-Lösung (Thermo Fisher Scientific, Waltham, USA). The reaction was stopped after 6 min by adding 6 % Orthophosphoric acid. Optical densities were measured using a plate spectrophotometer at 450 nm with a reference wavelength of 620 nm.

**Bulk RNA sequencing of human ileal biopsies**

Tissue preparation: Ileal Biopsies from patients were snap frozen in liquid nitrogen directly after the surgical procedure and stored at –80 °C until RNA extraction.

RNA extraction: For the bulk RNA sequencing analysis of human ileal biopsies, total RNA was isolated using a RNeasy plus Mini Kit (Quiagen, Netherlands) according to the manufacturer's recommendations. Measurement of RNA concentration was performed with a NanoDrop 2000 (Thermo Fisher Scientific, Waltham, USA). First, 1 µl RNase free water was measured as a blank. Then, 1 µl of each sample was measured to determine RNA concentration and purity.

Bulk mRNA sequencing and data analysis: The Illumina RNA sequencing analysis was performed by Novogene. In brief, sequencing libraries were generated using NEBNext® Ultra TM RNA Library Prep Kit for Illumina® (NEB, USA) and sequenced on an Illumina platform. Raw data (raw reads) of FASTQ format were processed through fastp. Mapping of the processed data to the reference genome homo sapiens (GRCh38/hg38) was performed using hisat2 (10). FeatureCounts was used for the quantification of the mapped reads (11). Raw mapped reads were processed in R (R version 4.3.3, Lucent Technologies) with DESeq2 (12), to determine differentially expressed genes and generate normalized read counts. Before processing the intestinal counts with DESeq2, raw counts were filtered for genes with zero transcripts. For PCA normalized counts were log transformed. The top 200 genes with most variable genes of PC1 were used for pathway enrichment analysis using the clusterProfiler package (maxGSsize = 500, pAdjustMethod = Benjamini-Hochberg, Ontology = Cellular component) (13). Heatmaps were generated using the pheatmap package (14). DEGs were defined as genes with a p-value $< 0.05$ and log2 Fold change (FC) $> |1.5|$ compared to controls and used for pathway enrichment analysis. Venn diagrams of DEGs were generated using the ggvenn package (15). Volcano plots were generated using the EnhancedVolcano package (16). Graphs were generated using the ggplot2 package (17) Deconvolution was performed with the MCPcounter package using the MCPcounter.estimate function with TPM converted counts and the reference genome GRCh38/hg38 (18, 19).
