## Supplementary Figures for "Microbiota-specific serum IgG links gut and joints through immune–endothelial crosstalk in arthritis"

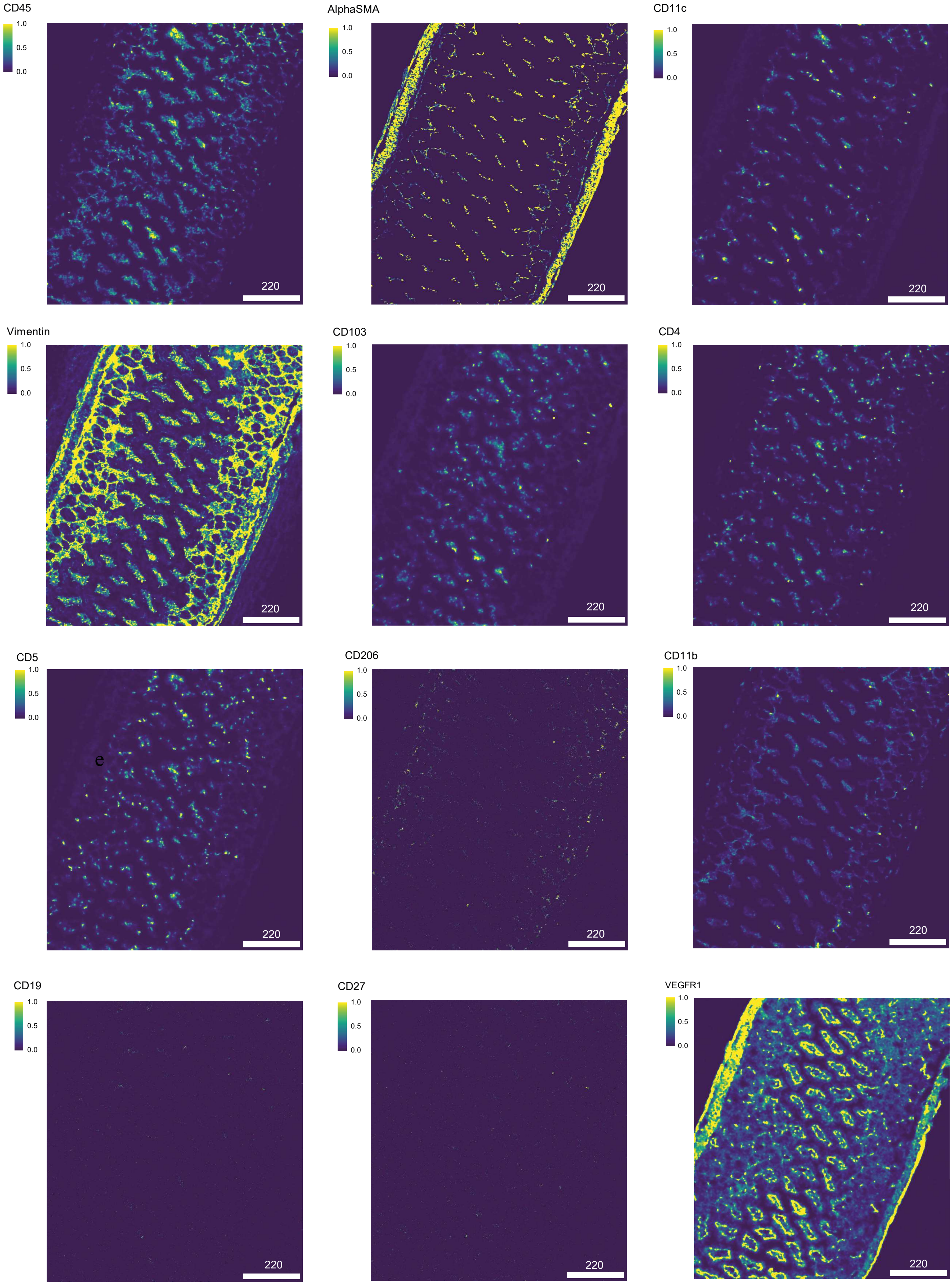
CD45
1.0
0.5
0.0
220
AlphaSMA
1.0
0.5
0.0
220
CD11c
1.0
0.5
0.0
220
Vimentin
1.0
0.5
0.0
220
CD103
1.0
0.5
0.0
220
CD4
1.0
0.5
0.0
220
CD5
1.0
0.5
0.0
e
220
CD206
1.0
0.5
0.0
220
CD11b
1.0
0.5
0.0
220
CD19
1.0
0.5
0.0
220
CD27
1.0
0.5
0.0
220
VEGFR1
1.0
0.5
0.0
220

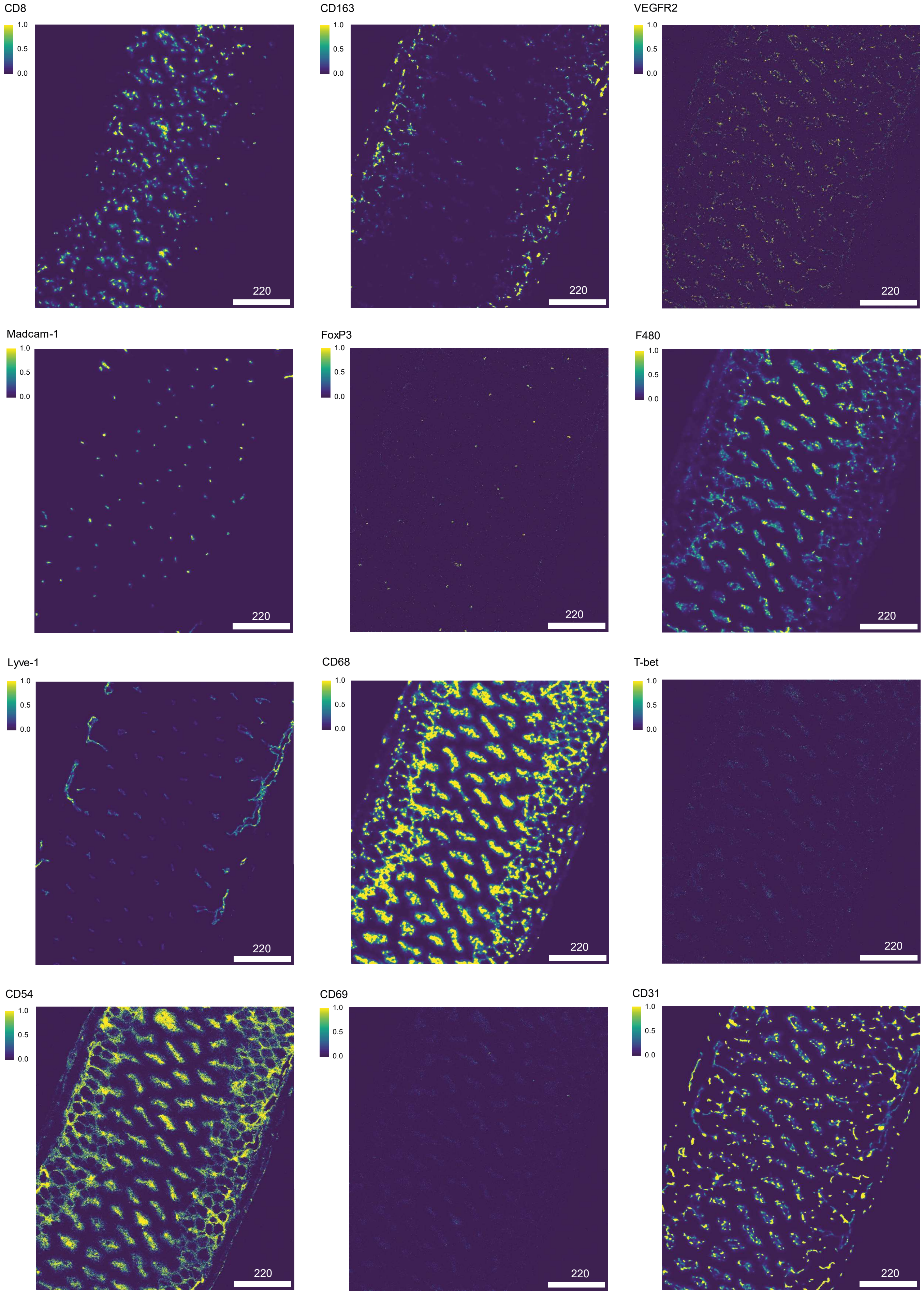
CD8
CD163
VEGFR2
Madcam-1
FoxP3
F480
Lyve-1
CD68
T-bet
CD54
CD69
CD31
1.0
0.5
0.0
220

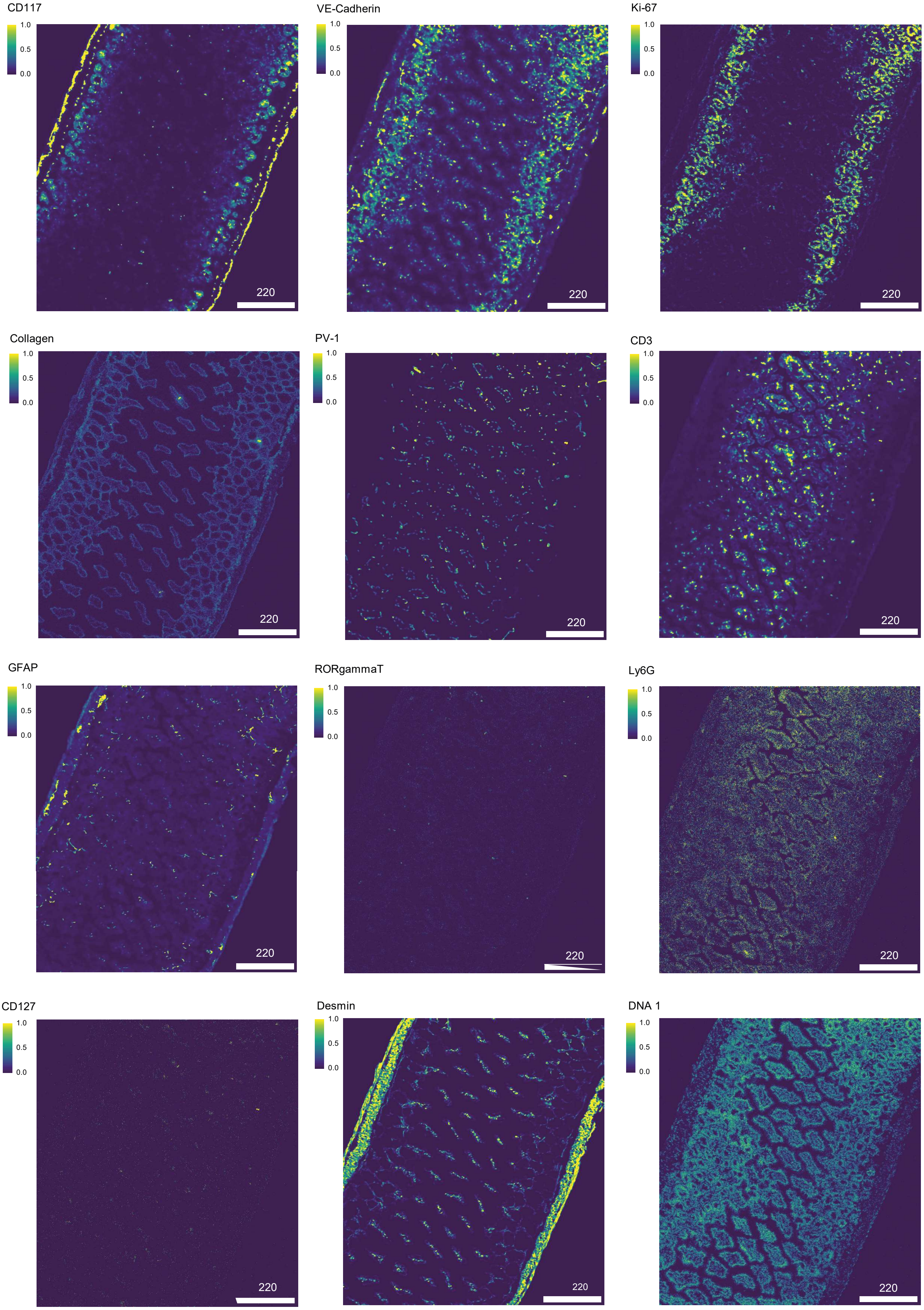
CD117
VE-Cadherin
Ki-67
Collagen
PV-1
CD3
GFAP
RORgammaT
Ly6G
CD127
Desmin
DNA 1
1.0
0.5
0.0
220

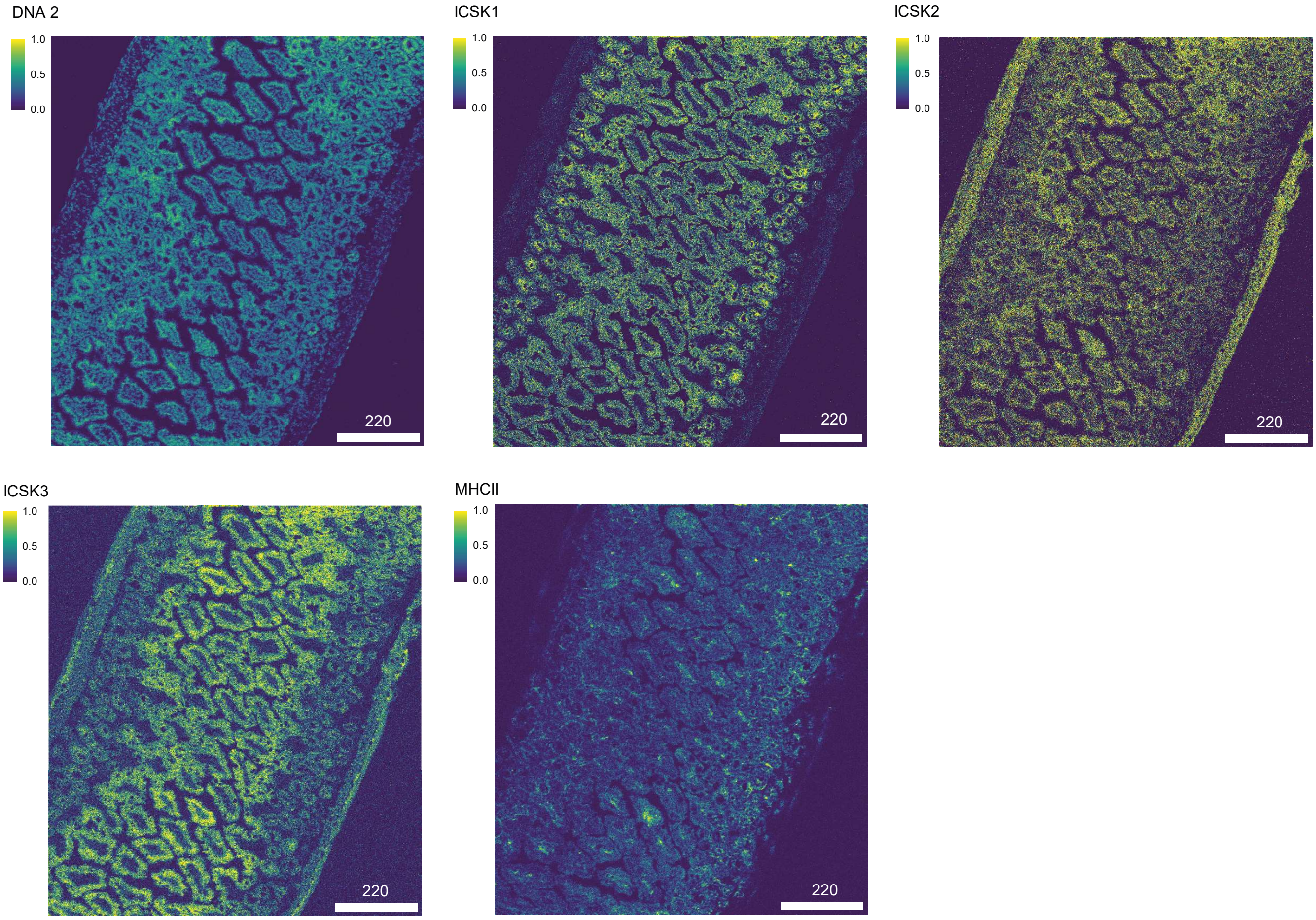


**Supplementary Fig. 1. Single marker expression of murine IMC panel.** Representative example ROI showing the normalized metal labelled marker expression of all used antibodies.

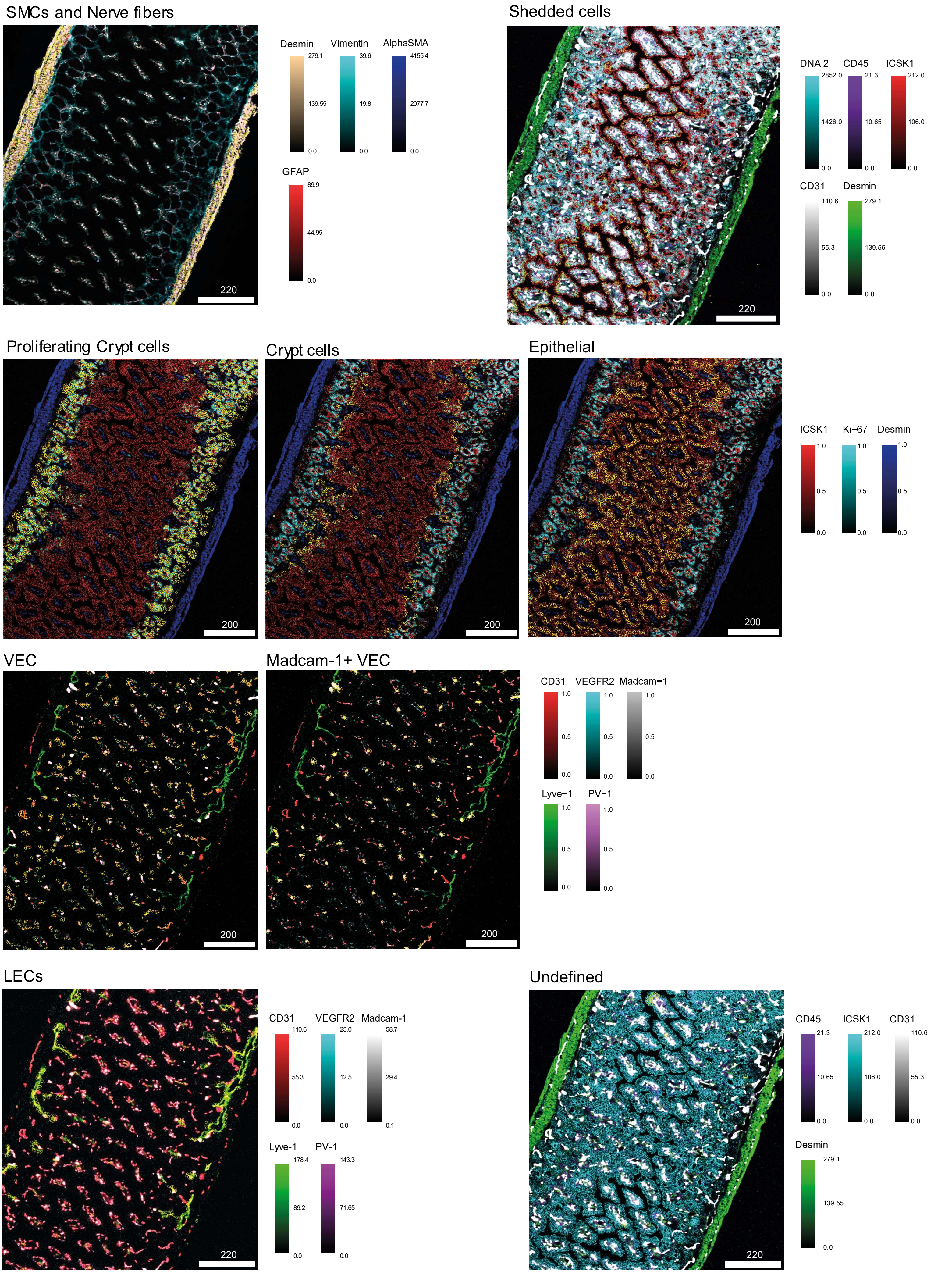

SMCs and Nerve fibers
Desmin
279.1
139.55
0.0
Vimentin
39.6
19.8
0.0
AlphaSMA
4155.4
2077.7
0.0
GFAP
89.9
44.95
0.0
220
Shedded cells
DNA 2
2852.0
1426.0
0.0
CD45
21.3
10.65
0.0
ICSK1
212.0
106.0
0.0
CD31
110.6
55.3
0.0
Desmin
279.1
139.55
0.0
220
Proliferating Crypt cells
200
Crypt cells
200
Epithelial
200
ICSK1
Ki−67
Desmin
1.0
0.5
0.0
VEC
200
Madcam-1+ VEC
200
CD31
VEGFR2
Madcam−1
1.0
0.5
0.0
Lyve−1
PV−1
1.0
0.5
0.0
LECs
220
CD31
110.6
55.3
0.0
VEGFR2
25.0
12.5
0.0
Madcam-1
58.7
29.4
0.1
Lyve-1
178.4
89.2
0.0
PV-1
143.3
71.65
0.0
Undefined
220
CD45
21.3
10.65
0.0
ICSK1
212.0
106.0
0.0
CD31
110.6
55.3
0.0
Desmin
279.1
139.55
0.0

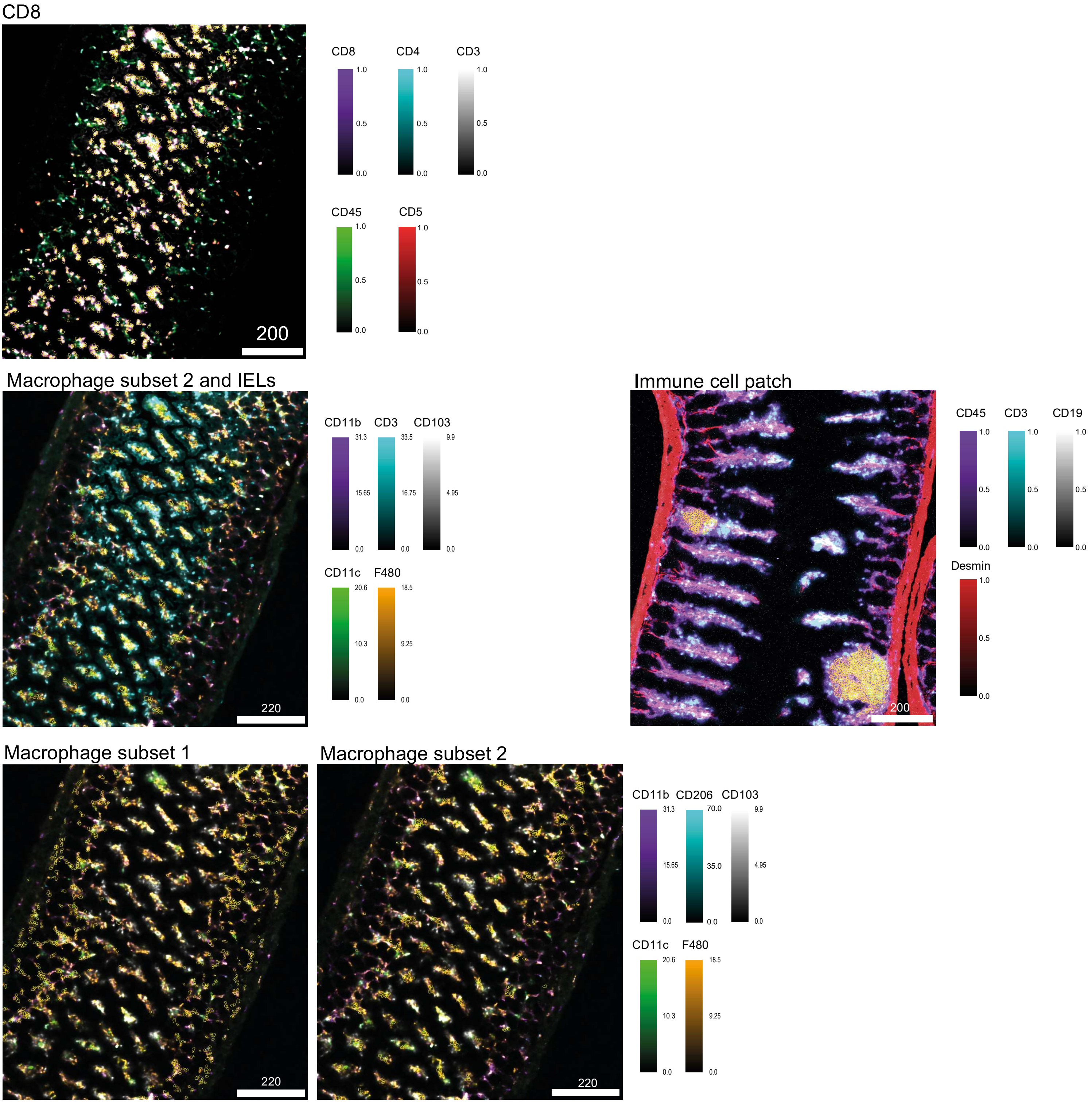


**Supplementary Fig. 2. With murine IMC panel identified cell types.** Representative example ROI showing the defined cell types (encircled in yellow) and the selected marker expression of the characteristic cell types.

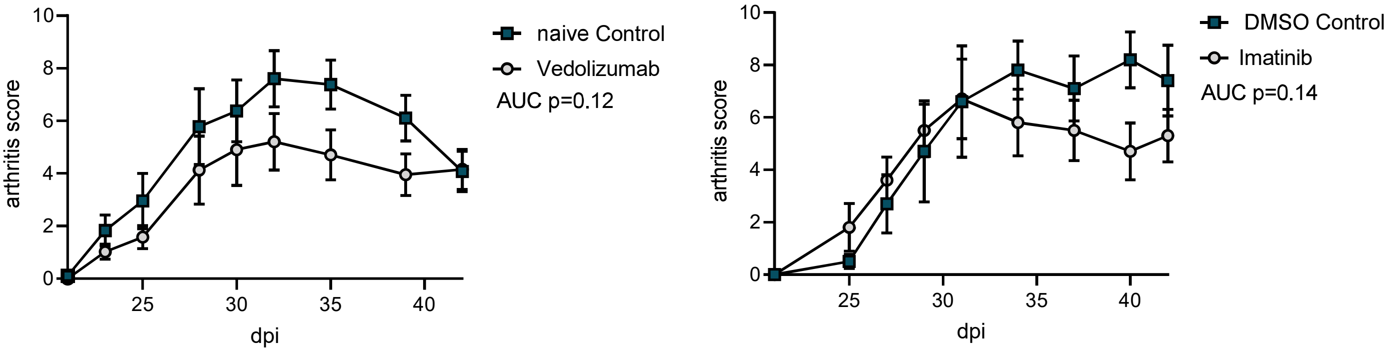


**Supplementary Fig. 3. Arthritis scores of CIA mice treated with a4b7 or Imatinib-mesylate compared to non-treated mice.** (**A**) Vedolizumab (α4β7 antibody) was administered by intraperitoneal injections (5 mg/kg bodyweight) 3 times per week starting 0 dpi until 21 dpi. (**B**) Imatinib-mesylate (Merck, Darmstadt, Germany) was solved in DMSO (100 mg/ml) and diluted in PBS for daily gavage from 15 dpi until 25 dpi (50 mg/kg bodyweight). Statistical difference between the groups was determined by unpaired t-test of AUC (area under the curve). $n \geq 5$.

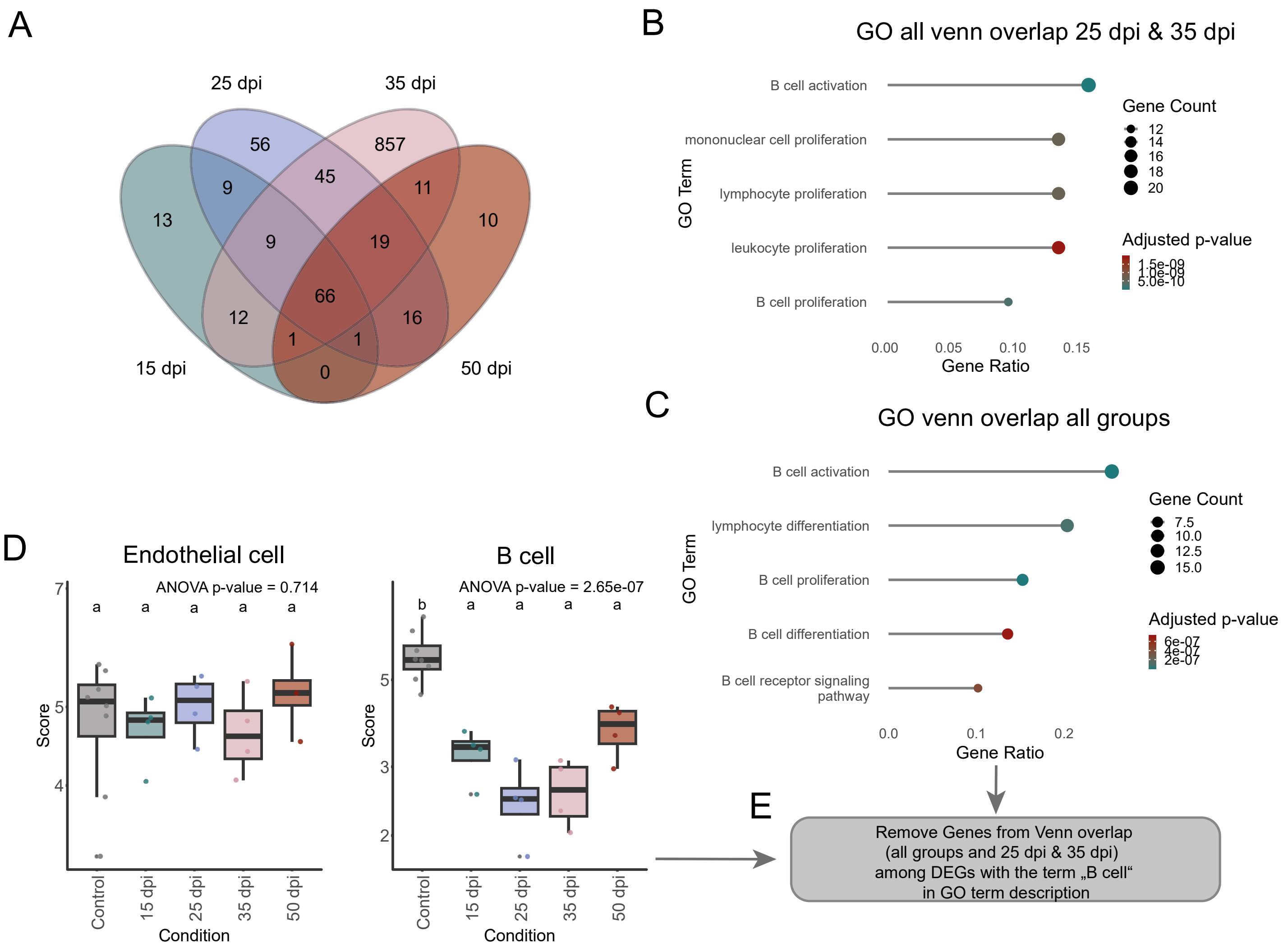


**Supplementary Fig. 4. Transcriptional signature of sorted bone CD31$^{+}$ cells reveals B-cell contamination. (A)** Venn diagram of differentially expressed genes (DEGs: padj < 0.05, log2FC >1.5 OR < -1.5). (B) GO analysis of DEGs from Venn overlap of all groups. (C) GO analysis of DEGs from Venn overlap of 26 dpi and 35 dpi. (D) Boxplot of estimated cell type scores (Endothelial cells and B cells) from deconvolution of bulk sequencing data using mMCP-Counter. Data are presented as boxplots showing the median (line), IGR (box), and whiskers extending to 1.5× IQR. Outliers are shown as individual points. Statistical analysis was performed using one-way ANOVA. For multiple comparisons between the groups Tukey-test was performed b p-value < 0.05 (E) Procedure to remove effect of B-cell contamination in samples. Differential expression analysis was performed using the DESeq2 package, with significance assessed based on p-values adjusted for multiple testing using the Benjamini-Hochberg FDR control method. (n ≥ 3)

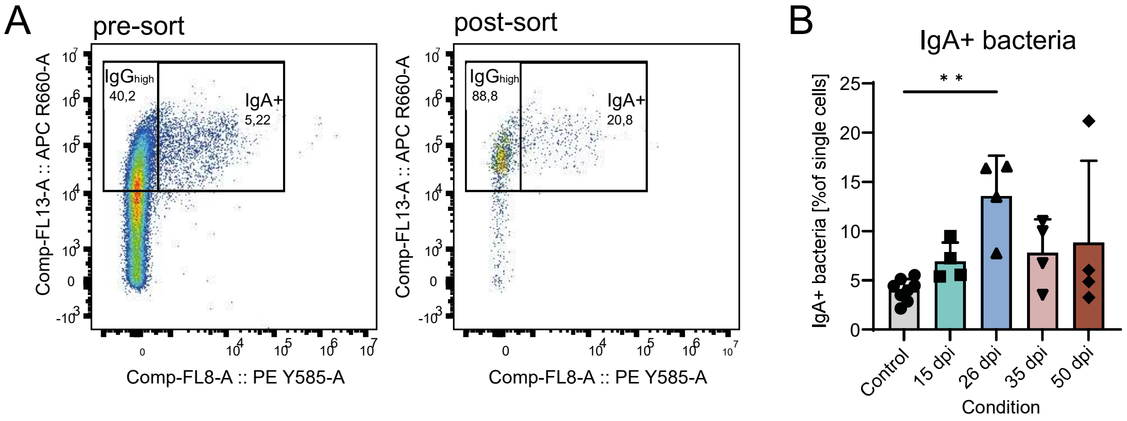


**Supplementary Fig. 5. FACS sorting of IgG$^{high}$ bacteria.** (**A**) Representative scatter plots (FACS plots) of sorted bacteria (pre-sort) and the reanalysis (post-sort). Cells were sorted for serum IgG$^{high}$ (APC) events. IgA positive gate (PE) was only used for quantification but not used for sorting. (**B**) Frequency of IgA coated bacteria at the different time points after immunization and naïve control mice. Statistical significance was assessed using one-way ANOVA. Multiple comparisons were performed using Tukey HSD. ($n \geq 4$) * p-value < 0.05

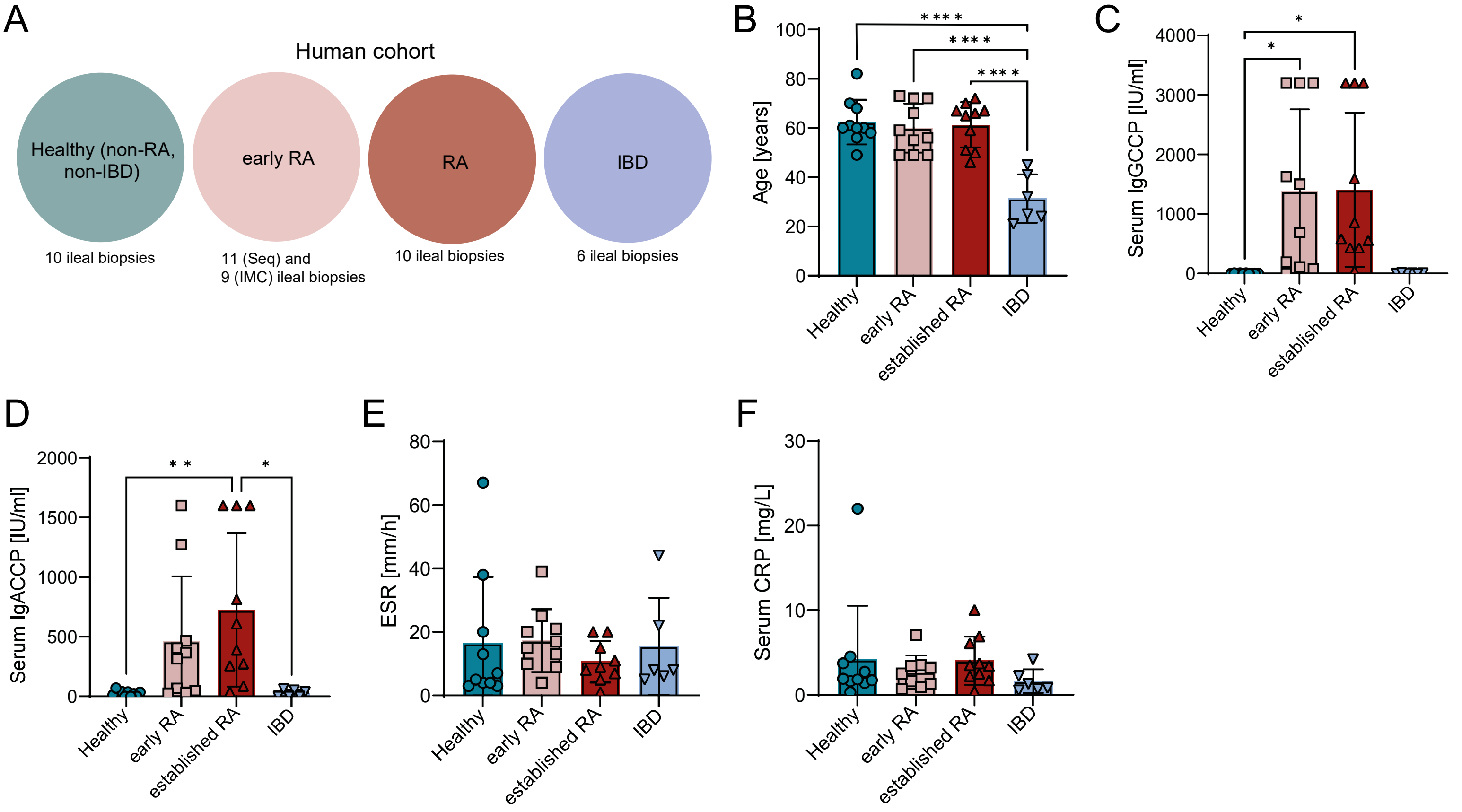


**Supplementary Fig. 6. Clinical parameters of human cohort. (A)** Graphical illustration of study cohort and analyzed samples. (**B**) Age of analyzed subjects in years at the time of the ileal biopsy sampling. (**C**) Serum IgGCCP serum levels in IU/ml at the time of the ileal biopsy sampling. (**D**) Serum IgACCP serum levels in IU/ml at the time of the ileal biopsy sampling. (**E**) Erythrocyte sedimentation rate (ESR) in mm/h at the time of the ileal biopsy sampling. (**F**) Serum C-reactive protein (CRP) levels in mg/l at the time of the ileal biopsy Data are presented as mean ± SD. Statistical significance was assessed using one-way ANOVA. Multiple comparisons were performed using Tukey HSD. ($n \geq 6$) * p-value < 0.05 ** p-value < 0.01 **** p-value < 0.0001

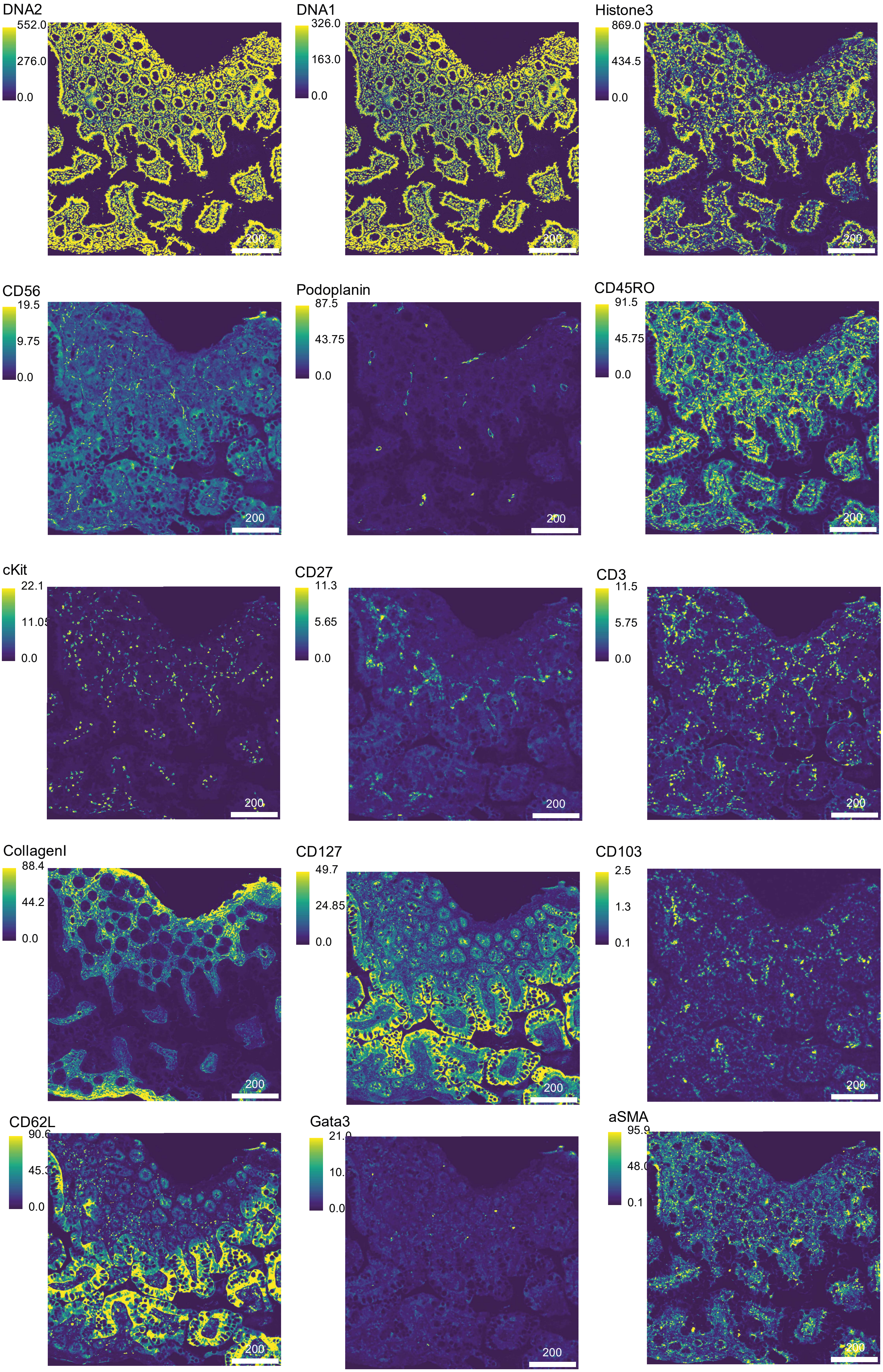
DNA2
552.0
276.0
0.0
DNA1
326.0
163.0
0.0
Histone3
869.0
434.5
0.0
CD56
19.5
9.75
0.0
Podoplanin
87.5
43.75
0.0
CD45RO
91.5
45.75
0.0
cKit
22.1
0.0
CD27
11.3
5.65
0.0
CD3
11.5
5.75
0.0
CollagenI
88.4
44.2
0.0
CD127
49.7
24.85
0.0
CD103
2.5
1.3
0.1
CD62L
0.0
Gata3
aSMA
0.1
200

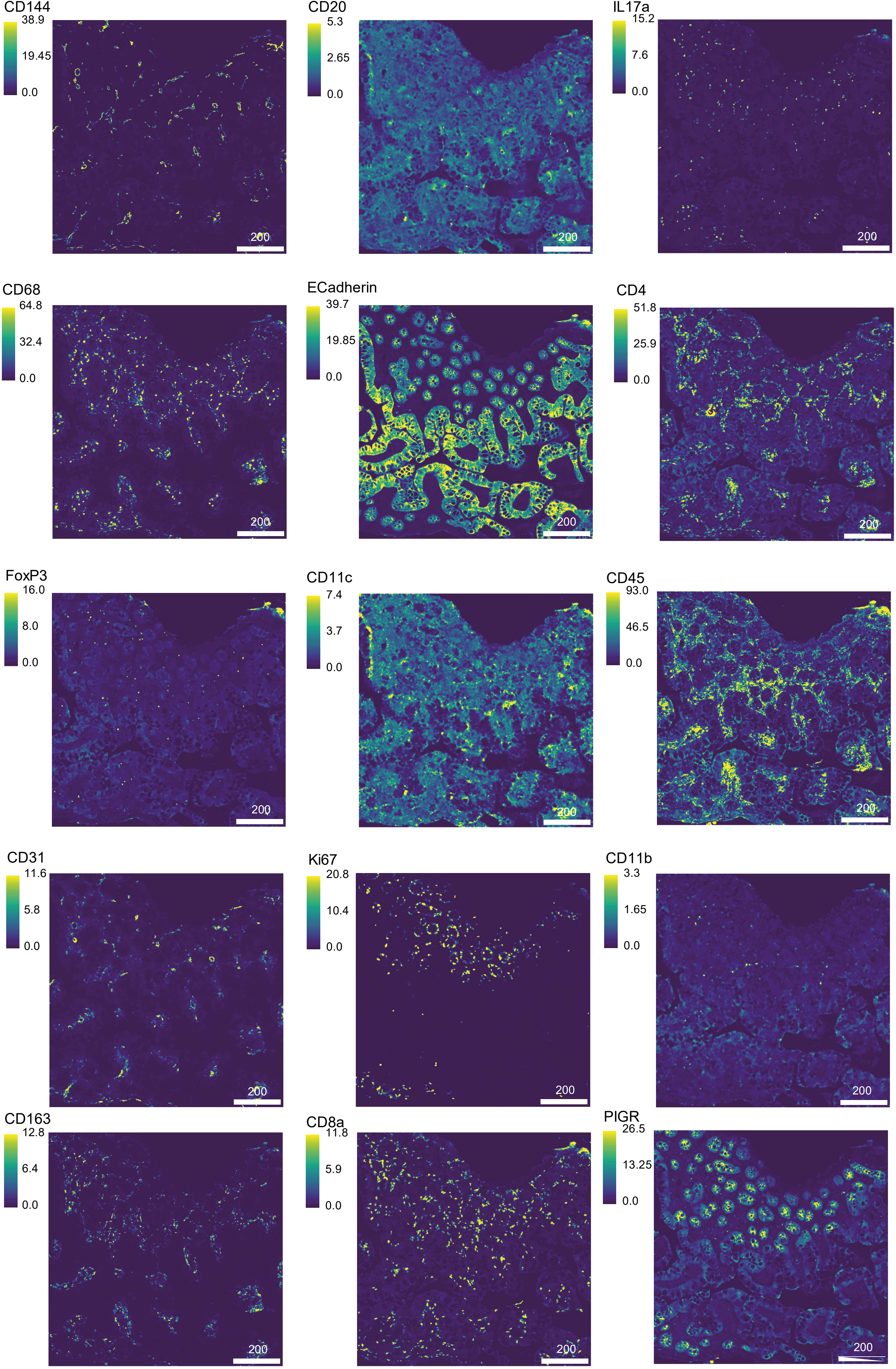
CD144
38.9
19.45
0.0
200
CD20
5.3
2.65
0.0
200
IL17a
15.2
7.6
0.0
200
CD68
64.8
32.4
0.0
200
ECadherin
39.7
19.85
0.0
200
CD4
51.8
25.9
0.0
200
FoxP3
16.0
8.0
0.0
200
CD11c
7.4
3.7
0.0
200
CD45
93.0
46.5
0.0
200
CD31
11.6
5.8
0.0
200
Ki67
20.8
10.4
0.0
200
CD11b
3.3
1.65
0.0
200
CD163
12.8
6.4
0.0
200
CD8a
11.8
5.9
0.0
200
PIGR
26.5
13.25
0.0
200

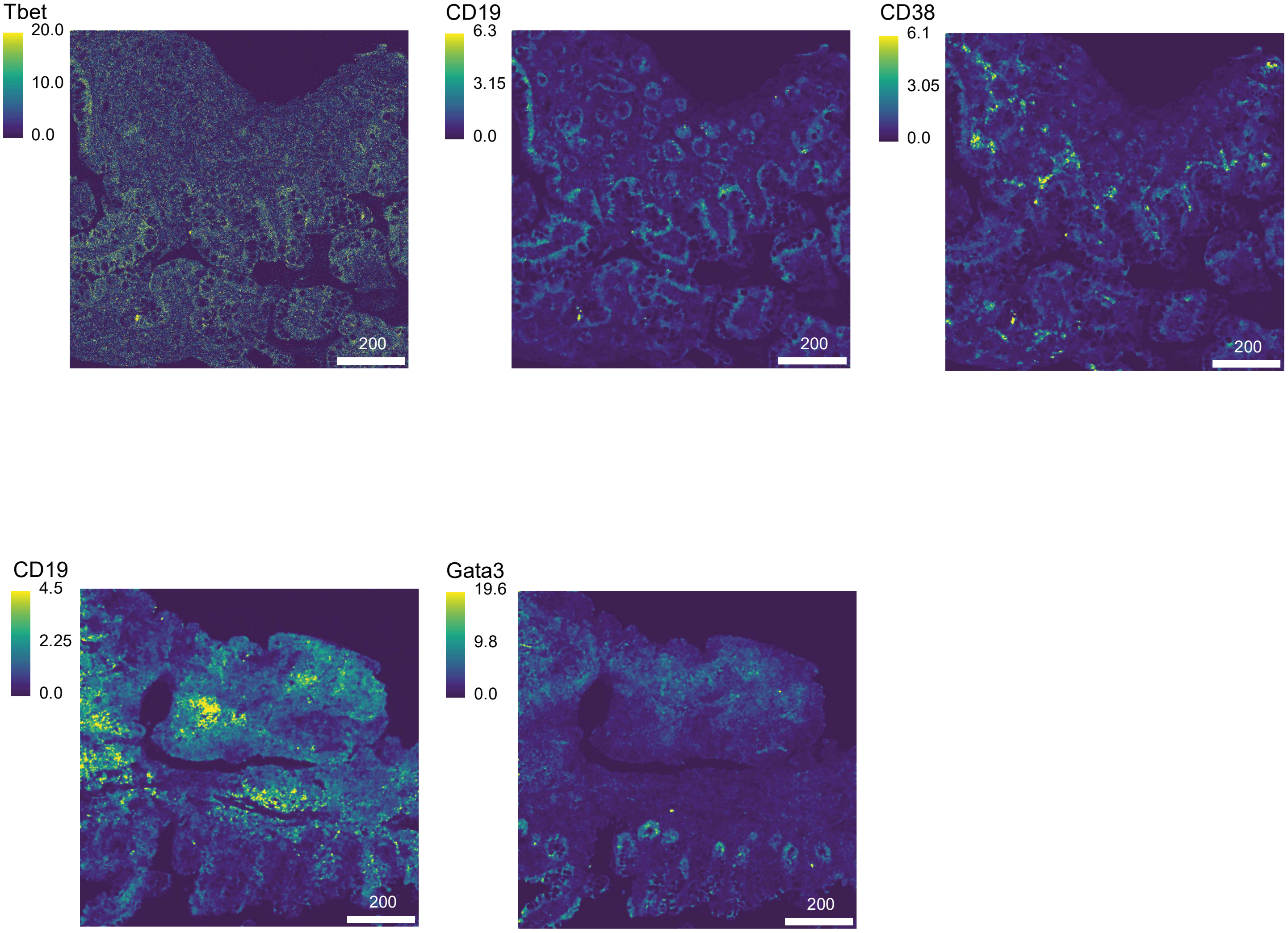


**Supplementary Fig. 7. Single marker expression of human IMC panel.** Representative example ROI showing the normalized metal labelled marker expression of all used antibodies.

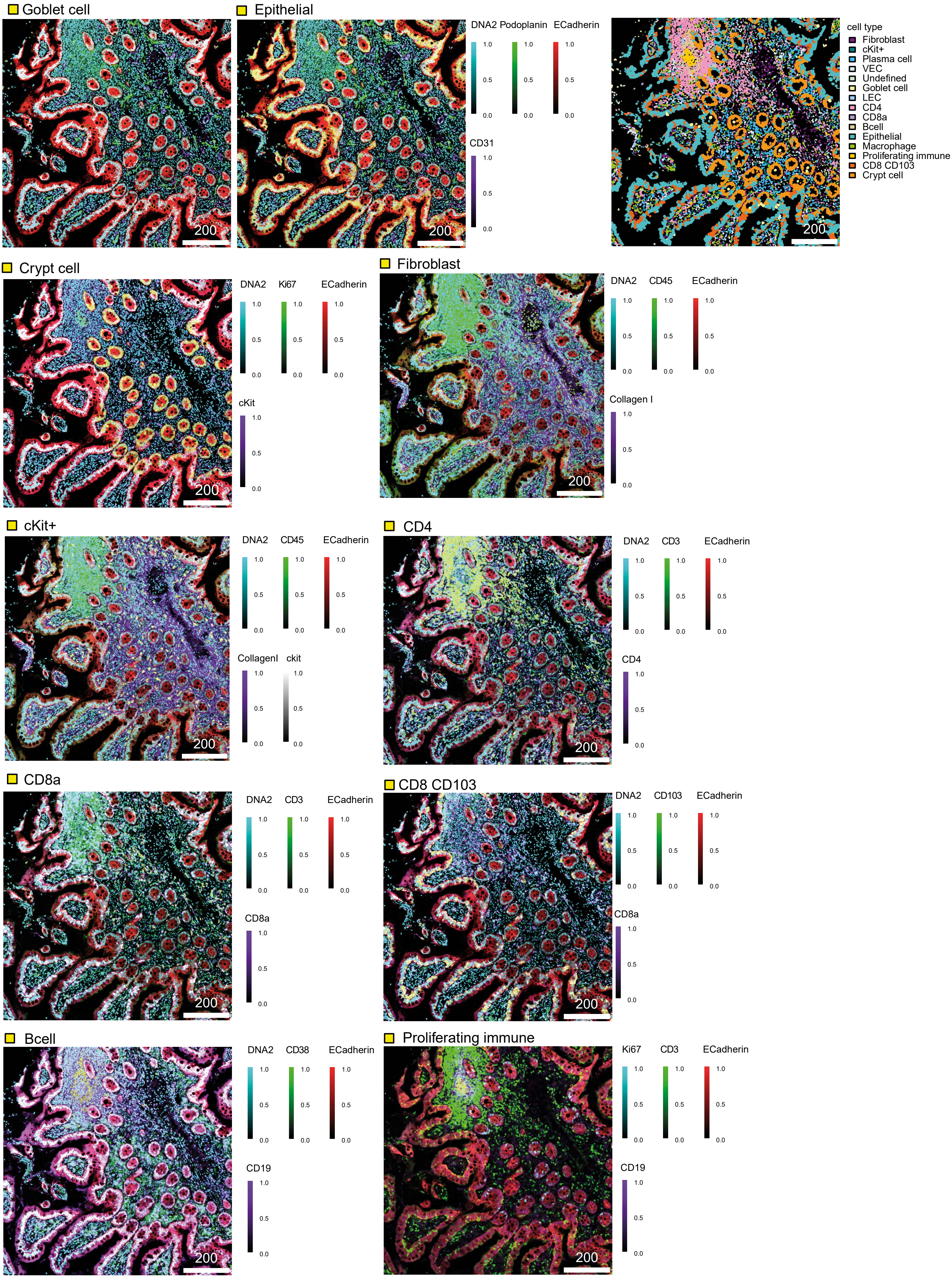

Goblet cell
200
Epithelial
200
DNA2 Podoplanin ECadherin
1.0
0.5
0.0
CD31
200
cell type
Fibroblast
cKit+
Plasma cell
VEC
Undefined
Goblet cell
LEC
CD4
CD8a
Bcell
Epithelial
Macrophage
Proliferating immune
CD8 CD103
Crypt cell
Crypt cell
200
DNA2 Ki67 ECadherin
cKit
Fibroblast
200
DNA2 CD45 ECadherin
Collagen I
cKit+
200
DNA2 CD45 ECadherin
CollagenI ckit
CD4
200
DNA2 CD3 ECadherin
CD4
CD8a
200
DNA2 CD3 ECadherin
CD8a
CD8 CD103
200
DNA2 CD103 ECadherin
CD8a
Bcell
200
DNA2 CD38 ECadherin
CD19
Proliferating immune
200
Ki67 CD3 ECadherin
CD19

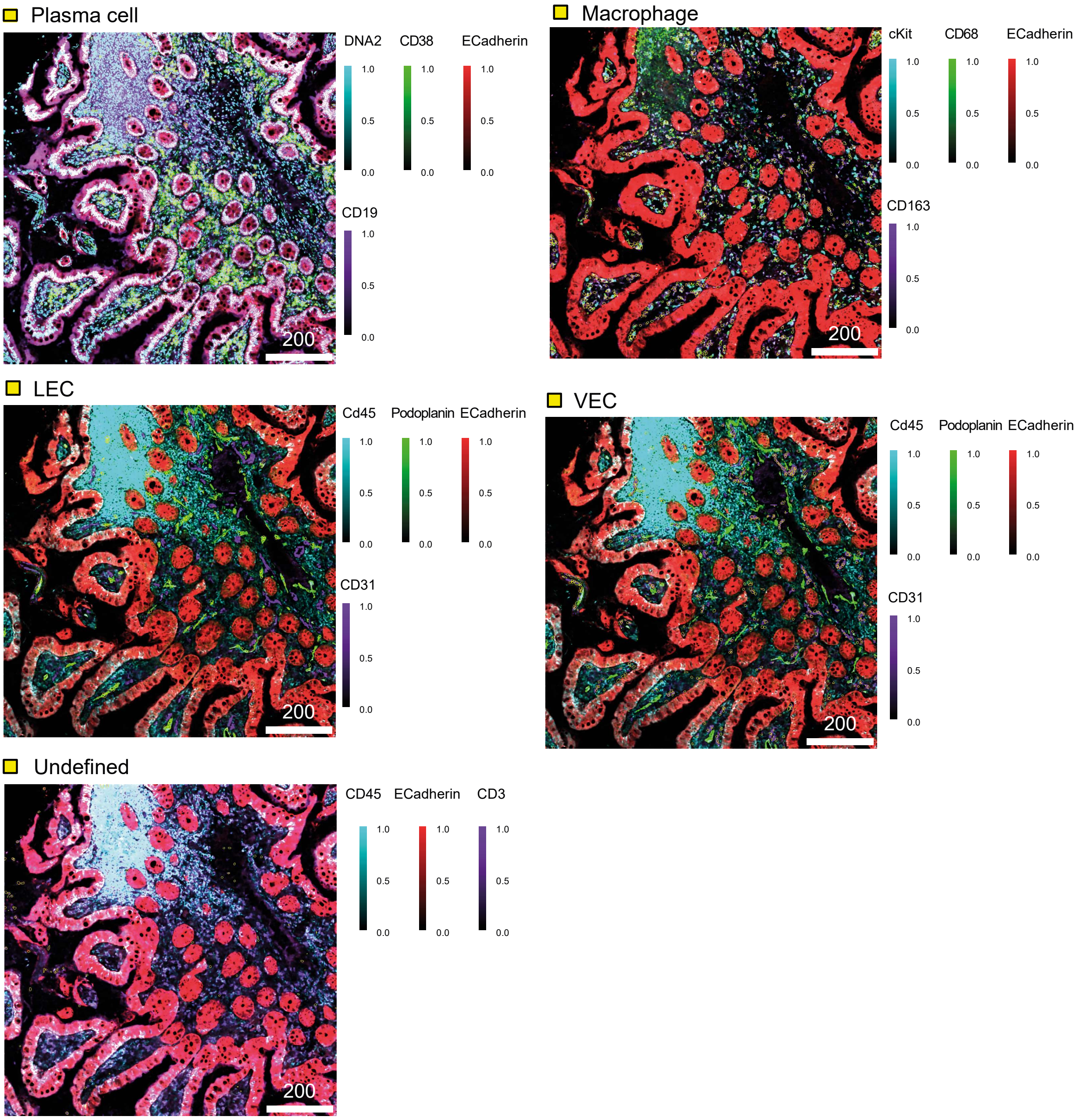


**Supplementary Fig. 8. With human IMC panel identified cell types.** Representative example ROI showing the defined cell types (encircled in yellow) and the selected marker expression of the characteristic cell types.

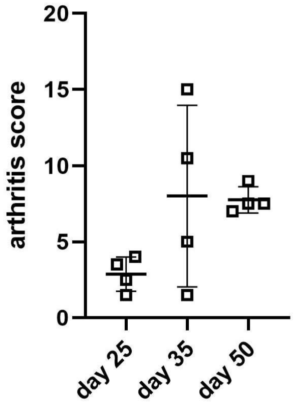


**Supplementary Fig. 8. Arthritis scores of mice used for intestinal endothelial cell RNA-Seq. after the second immunization** 25, 35 or 50 days post-immunization (dpi). Scores were measured on the respective sacrifice date. n = 4 mice per time point.

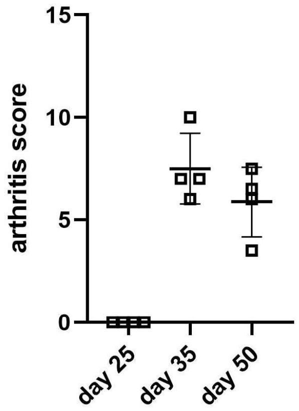


**Supplementary Fig. 9. Arthritis scores of mice used for bone endothelial cell RNA-Seq. after the second immunization** 25, 35 or 50 days post-immunization (dpi). Scores were measured on the respective sacrifice date. n = 4 mice per time point.
